# Cell-type, single-cell, and spatial signatures of brain-region specific splicing in postnatal development

**DOI:** 10.1101/2020.08.27.268730

**Authors:** Anoushka Joglekar, Andrey Prjibelski, Ahmed Mahfouz, Paul Collier, Susan Lin, Anna Katharina Schlusche, Jordan Marrocco, Stephen R. Williams, Bettina Haase, Ashley Hayes, Jennifer G. Chew, Neil I Weisenfeld, Man Ying Wong, Alexander N. Stein, Simon Hardwick, Toby Hunt, Zachary Bent, Olivier Fedrigo, Steven A. Sloan, Davide Risso, Erich D. Jarvis, Paul Flicek, Wenjie Luo, Geoffrey S. Pitt, Adam Frankish, August B. Smit, M. Elizabeth Ross, Hagen U. Tilgner

## Abstract

Alternative RNA splicing varies across brain regions, but the single-cell resolution of such regional variation is unknown. Here we present the first single-cell investigation of differential isoform expression (DIE) between brain regions, by performing single cell long-read transcriptome sequencing in the mouse hippocampus and prefrontal cortex in 45 cell types at postnatal day 7 (www.isoformAtlas.com). Using isoform tests for brain-region specific DIE, which outperform exon-based tests, we detect hundreds of brain-region specific DIE events traceable to specific cell-types. Many DIE events correspond to functionally distinct protein isoforms, some with just a 6-nucleotide exon variant. In most instances, one cell type is responsible for brain-region specific DIE. Cell types indigenous to only one anatomic structure display distinctive DIE, where for example, the choroid plexus epithelium manifest unique transcription start sites. However, for some genes, multiple cell-types are responsible for DIE in bulk data, indicating that regional identity can, although less frequently, override cell-type specificity. We validated our findings with spatial transcriptomics and long-read sequencing, yielding the first spatially resolved splicing map in the postnatal mouse brain (www.isoformAtlas.com). Our methods are highly generalizable. They provide a robust means of quantifying isoform expression with cell-type and spatial resolution, and reveal how the brain integrates molecular and cellular complexity to serve function.

## Introduction

Alternative splicing (AS) affects almost all spliced genes in mammals^1,2^, vastly expands the proteome^3^ and increases functional diversity of cell types^4^. Alternative transcription start sites (TSS) and poly-adenylation (polyA) sites further expand the alternative isoform landscape, regulating development, differentiation, and disease^5–9^. These RNA variables often depend on each other^10–13^, and how their combined status impacts individual molecules can only be assessed using long-read sequencing^11,12,14–17^, which sequences transcripts in single reads with no assembly required, thereby reducing alternative transcript assembly errors and enabling accurate isoform quantification.

Brain AS is especially diverse^18^ and brain-region specific expression patterns of splicing factors^19^ and other RNA-binding proteins^20^ drive brain-region specific splicing. Examples include diseases implicated genes such as *MAPT, Bin1*, and neurexins^16,17,21^. Brain-region specific isoform expression can either originate from molecular regulation in one or multiple cell types, or can arise purely from gene-expression or celltype abundance differences without splicing regulation. These distinct models are especially important during postnatal development. For instance, in hippocampus (HIPP) and prefrontal cortex (PFC), multiple cell types undergo differentiation, which is influenced by development-specific splicing^1,22–25^ distinct from that of mature cell types. However, no cell-type specific isoform investigation across brain regions exists to-date, owing to limitations in technology, throughput, and testing methods. HIPP and PFC are highly specialized regions of the telencephalon, and their circuitry is heavily implicated in movement control, cognition, learning, and memory formation. Disorders involving HIPP and PFC manifest in cognitive deficits, and understanding changes occurring at crucial developmental timepoints of these structures is important for case-control studies. Here, we employ single-cell isoform RNA sequencing (ScISOrSeq)^26^ with increased throughput in HIPP and PFC at mouse postnatal day 7 (P7) to test and define cell-type specific contributions to brain-region specific splicing. Furthermore, we devised a spatial isoform expression method, which provides a spatial exon expression map (see www.isoformAtlas.com) in addition to the existing spatial gene-expression map of the Allen developing brain atlas.

## Results

### Short read clustering of P7 hippocampus and prefrontal cortex tissue assigns precursors to known adult cell-types

Our ScISOrSeq approach used barcoded single cells followed by both short and long-read analyses to reveal splice variants specific to cell types (Fig 1a). We identified cell types first using single-cell 3’seq sequencing. Short-read clustering across two hippocampal replicates revealed no need for integration anchors^27^ to correct for batch effects (Fig S1a). Characteristic markers^28,29^ for 24 clusters in HIPP (Fig S1b) identified eight glial types, including two astrocyte, three oligodendrocyte, a radial glia like (RGLs), ciliated ependymal, and secretory choroid plexus epithelial (CPE) cell clusters. Furthermore, we observed six non-glial, non-neuronal populations including vascular endothelial and microglia and macrophage myeloid cells (Fig 1b). RNA velocity analysis revealed neuronal lineages in various differentiation stages (Fig S1c-d): a neuronal intermediate progenitor cell (NIPC) population; three dentate gyrus granule neuroblast clusters (DG-GranuleNB); and three clusters each of excitatory (EN) and inhibitory neurons (IN). Alignment of our P7 data with published P30 hippocampal data^30^ revealed subtype identities (CA3, CA1, Subiculum) for three excitatory neuron clusters and medial ganglionic eminence (MGE) and non-MGE derived interneurons^31–33^ (*PV*+, *Sst*+, *Lamp5*+, *Vip*+) in one cluster (IN1), distinct from Cajal-Retzius (IN3) cells (Figs 1b, S1e-g).

**Figure 1 -.**
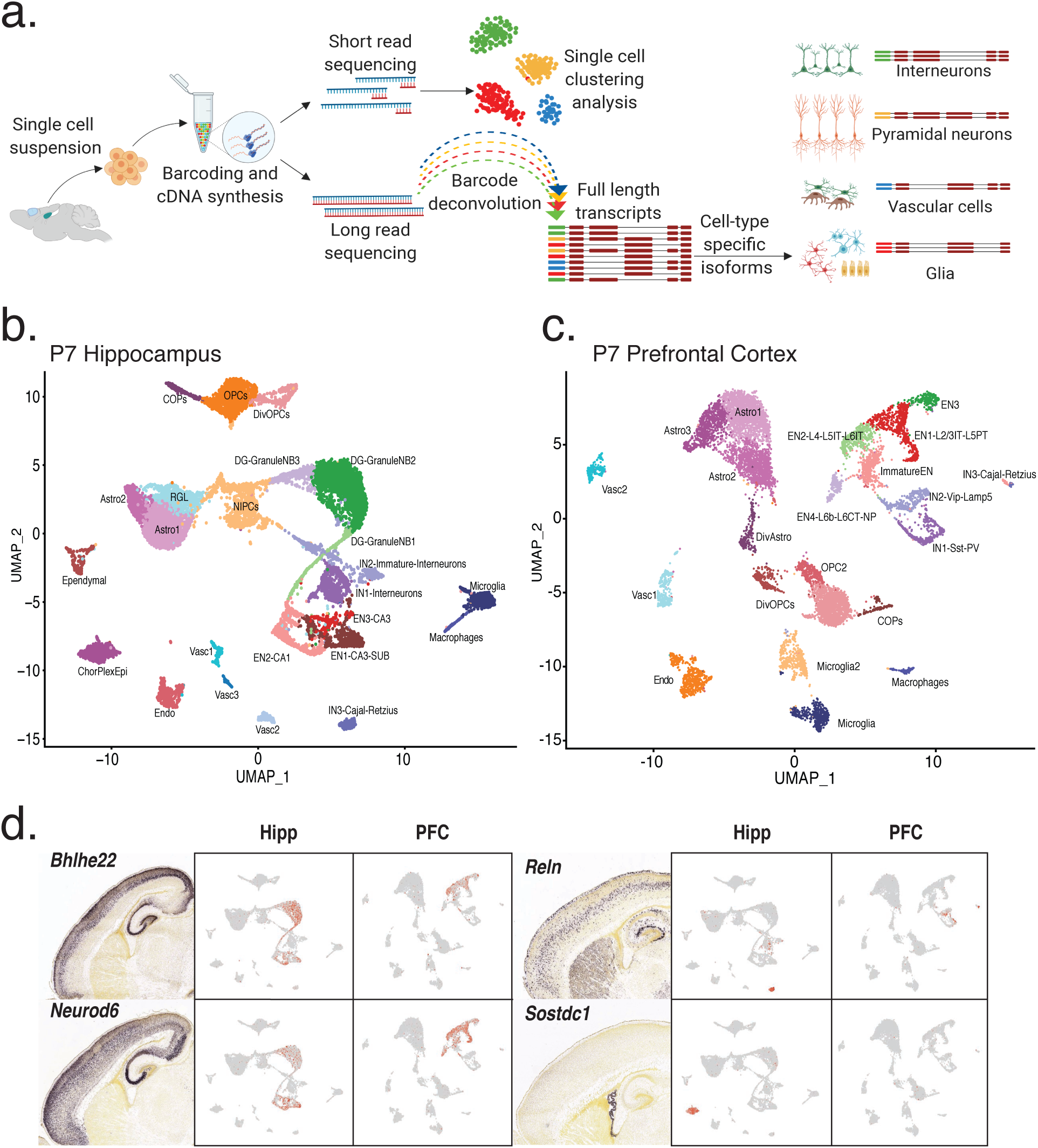
Short read clustering of P7 hippocampus and prefrontal cortex tissue recovers precursors to known adult cell-types. **a.** Schematic of the ScISOrSeq workflow (created with BioRender.com) **b.** UMAP of P7 hippocampus (HIPP) data. Cell-types identified by marker genes, RNA velocity analysis, and alignment to published data shown in S1 **c.** UMAP of P7 prefrontal cortex (PFC) data. Cell-types identified by marker genes, RNA velocity analysis, and alignment to published data shown in S2 **d.** In situ hybridization (ISH) images from Allen developing mouse brain atlas for marker genes and corresponding projections on UMAP plots from HIPP and PFC.

Similar analysis in PFC revealed seven glial clusters including astrocytes, oligodendrocytes, six populations of non-glial cells including vascular endothelial cells, myeloid cells, and seven neuronal types^34,35^ with confirmation of intermediate states from RNA velocity (Fig 1c, Fig S2a-d). Alignment with public P30 cortex data^30^ further subdivided neuronal clusters into known cortical excitatory and interneuron classes (Fig S2e-g). In contrast to HIPP, the MGE (*PV*+, *Sst*+) and non-MGE interneurons (*Vip*+, *Lamp5*+) in the PFC were better separated into two clusters (IN1, IN2), while Cajal-Retzius cells again clustered separately (IN3). We identified excitatory neurons corresponding to different cortical layers which are not well-differentiated at P7. P4 ISH images (Image credit: Allen Institute) alongside gene expression projected onto the UMAP plots further validated our cell-type identification for both regions (Fig 1d).

### A gene-wise test to determine differential isoform expression (DIE)

We next conducted long-read sequencing on our single-cell full-length HIPP and PFC cDNA (Supplementary Table 1), and deconvolved reads for each cell type using single-cell barcodes (Fig 1a) for two independent replicates (Figs S3, S4). Differential exon usage between two conditions has been successfully assessed using a 2×2 contingency table per exon^2^. Using this method on our long-read data from HIPP and PFC yielded 31 genes (1.45%, n=2132) exhibiting differential exon usage after Benjamini-Yekutieli (BY) correction^36^ for dependent tests (Fig S5a). Given this harsh correction, we devised a more sensitive gene-level test that considers TSS and polyA-sites in addition to exon connectivity. In this test, we count isoforms per gene in both conditions, leading to a nx2 table. This yields fewer and independent tests, allowing for the Benjamini-Hochberg (BH) correction^36^ and reducing false negatives. In each sample, we define “percent isoform” (Π) as an isoform’s relative abundance among its gene’s transcripts. Similarly to requiring a ΔΨ>=0.1 for short reads^2^, we require FDR<=0.05 and ΔΠ ≥ 0.1, contributed collectively by at most two isoforms to consider a gene exhibiting differential isoform expression (DIE, Fig 2a).

**Figure 2 -.**
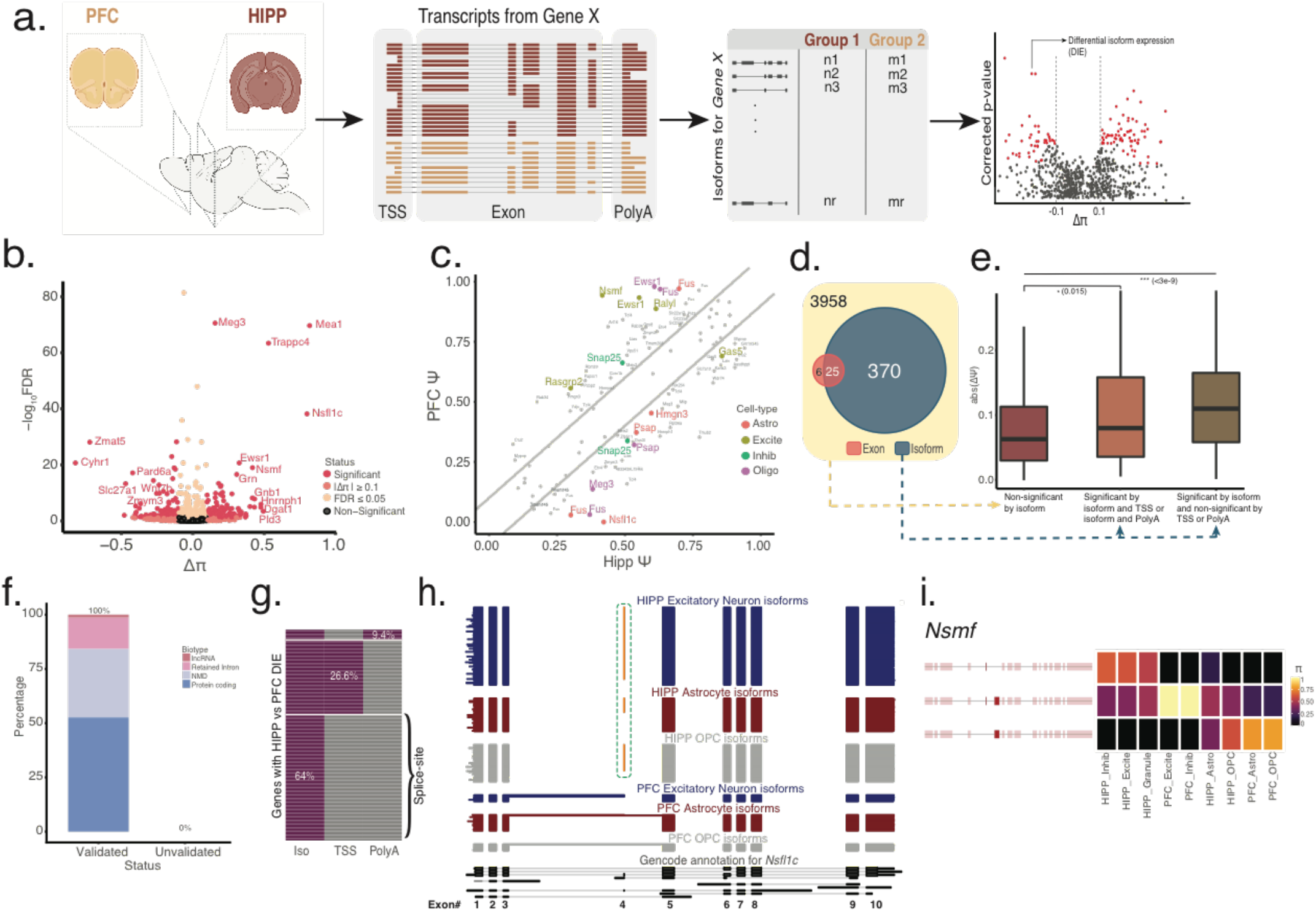
A gene-wise test for differential isoform expression (DIE) is more sensitive than an exon-wise test at detecting splicing changes. **a.** Schematic of the scisorseqR approach - Barcode deconvolution, filtering, pairwise comparison, and reporting of significant results based on FDR and Δπ cutoffs (created with BioRender.com) **b.** Volcano plot of bulk HIPP vs. bulk PFC differential abundance analysis, with the effect size (Δπ) on the X-axis and BH corrected p-value on the Y-axis. Points are colored according to the levels of significance based on FDR and Δπ value. Genes considered significant (pink) when FDR <= 0.05 and |Δπ| >= 0.1 **c.** Percentage of novel transcripts by scisorseqR that were manually validated as being novel by Gencode team, and breakdown of predicted function, **d.** Heatmap of significant DIE genes according to entire isoform between bulk HIPP and bulk PFC that also exhibit differential usage of transcription start site (TSS) and polyA-site (PolyA). Each row is a single gene. Grey represents genes that are non-significant by category (Iso/TSS/PolyA) whereas purple represents significant by category **e.** Scatter plot showing the Δψ of all exons for genes that are show significant DIE between HIPP and PFC. Grey points represent non-significant exons. Points are colored according to the cell-type in which an exon is considered significant by a BY corrected p-value and a Δπ>= 0.1. Diagonal lines indicate cutoff of 0.1 Δψ **f.** Venn diagram showing the overlap of genes significant by DIE and genes significant by exon tests **g.** Boxplot showing the maximum absolute value of Δψ per gene in three different categories: Genes that are not significant by DIE tests, genes that are significant by DIE tests and also exhibit differential TSS or polyA-site usage, and genes that are significant by DIE and do not exhibit differential TSS or polyA-site usage **h.** Isoform expression of NsfHc gene. Each horizontal line in the plot represents a single transcript colored according to the cell-type it is represented in. Therefore, blocks represent exons and whitespace represents intronic space (not drawn to scale). Orange exon represents alternative inclusion **i.** Isoform expression for the Nsmf gene. Each row represents an isoform colored by π and each column represents a cell-type in HIPP or PFC.

In contrast to the 31 significant genes derived by the exon-based test, 395 genes (FDR <= 0.05, ΔΠ >= 0.1; 9.06%; n=3958) exhibited DIE when comparing HIPP and PFC isoforms using the gene-level test (Fig 2b). The multiple testing correction factor influenced the significant gene number. For example, the *H13* gene (Fig S5b) had a p-value of 1.7×10^-4^ (uncorrected) and 2.7×10^-3^ (corrected) by isoform tests, with a Benjamini-Hochberg correction factor of 15.6. The same gene’s alternative exon had an p-value of 1.3×10^-4^ (uncorrected) and 0.057 (corrected, non-significant) by exon tests with a Bejamini-Yekutieli correction factor of 431.3. Thus, gene-wise isoform testing is more sensitive (Fig 2c-d). Concordantly, the maximum ΔΨ for genes with DIE is higher than genes without DIE (p≤ 0.015, Wilcoxon-rank-sum test) (Fig 2e).

Since our gene-level test considers exon connectivity, we can identify the exact isoforms contributing to DIE. Among the top two contributing isoforms for the 395 genes exhibiting regional DIE, we identified 76 high-confidence novel isoforms (Methods). Manual validation using GENCODE criteria confirmed all 76. Functionally, 40 (52.6%) are coding transcripts, 24 (31.6%) show nonsense mediated decay (NMD), 11 (14.5%) show intron retention, and one isoform is of a long noncoding gene (*Meg3*) (Fig 2f). Such noncoding and NMD transcripts indicate region specific regulation^1^. To pinpoint the source of isoform differences in the 395 genes, we tested TSS and polyA-site abundance per gene across brain regions, which follow the same statistical framework as the isoform tests. 141 (of 395) genes exhibited differential TSS or polyA-site usage (Fig 2g). By extension, the remaining 254 genes are explained by splice-site usage differences.

Many genes with DIE have ΔΠ≤ 0.5; however we also identified drastic switches (ΔΠ≥0.5): *Nsfl1c* encodes the Nsfl1 cofactor p47, which regulates tubular ER formation, influences neuronal dendritic spine formation and dendritic arborization^37,38^. A 6nt microexon is preferentially included in HIPP across neuronal and glial cell types but is absent in the same PFC cell types (Fig 2h). The synaptic gene *Nsmf*, encoding the NMDA receptor activated protein Jacob, is involved in the cAMP pathway^39,40^ and through nuclear translocation of Jacob in memory formation^39,40^. In our data, the major HIPP isoform is completely absent in PFC, while the second HIPP isoform represents the majority of that gene’s PFC expression. The isoforms differ by a 69nt exon with a nuclear localization signal and one of two synaptic targeting elements. Hence, this exon may affect the synapse to nucleus signaling that the protein is involved in. A third *Nsmf* isoform with the 69nt exon but lacking a 6nt micro-exon is favored in PFC over HIPP but completely missing from neuronal cells, highlighting the regulatory role of micro-exons in neuronal function^1,41^ (Fig 2i).

### Differential isoform expression across brain regions is governed predominantly by one specific cell type

Gene expression transcript similarities among clusters defined a cell-type hierarchy, first separated by neurons and non-neurons, and then by other cell types (Fig S6). Since inhibitory neuron types are transcriptionally more similar to each other than to excitatory types^34,35^, we grouped finer inhibitory neuron subtypes (IN1, IN2, IN3) into a composite inhibitory neuron (IN) category, and excitatory neuron subtypes into an excitatory neuron (EN) category. We hypothesized three alternative models that could underlie differential isoform expression between brain regions: Brain-region DIE could be contributed by 1) multiple or all cell types that change splice variants (‘Both-Cell-Types-Model’); 2) a single cell type that changes splice variants (‘Single-Cell-Type-Model’); or 3) by changes in expression or cell-type abundance without any change in splice variants (‘No-Cell-Type-Model’) (Fig 3a).

**Figure 3 -.**
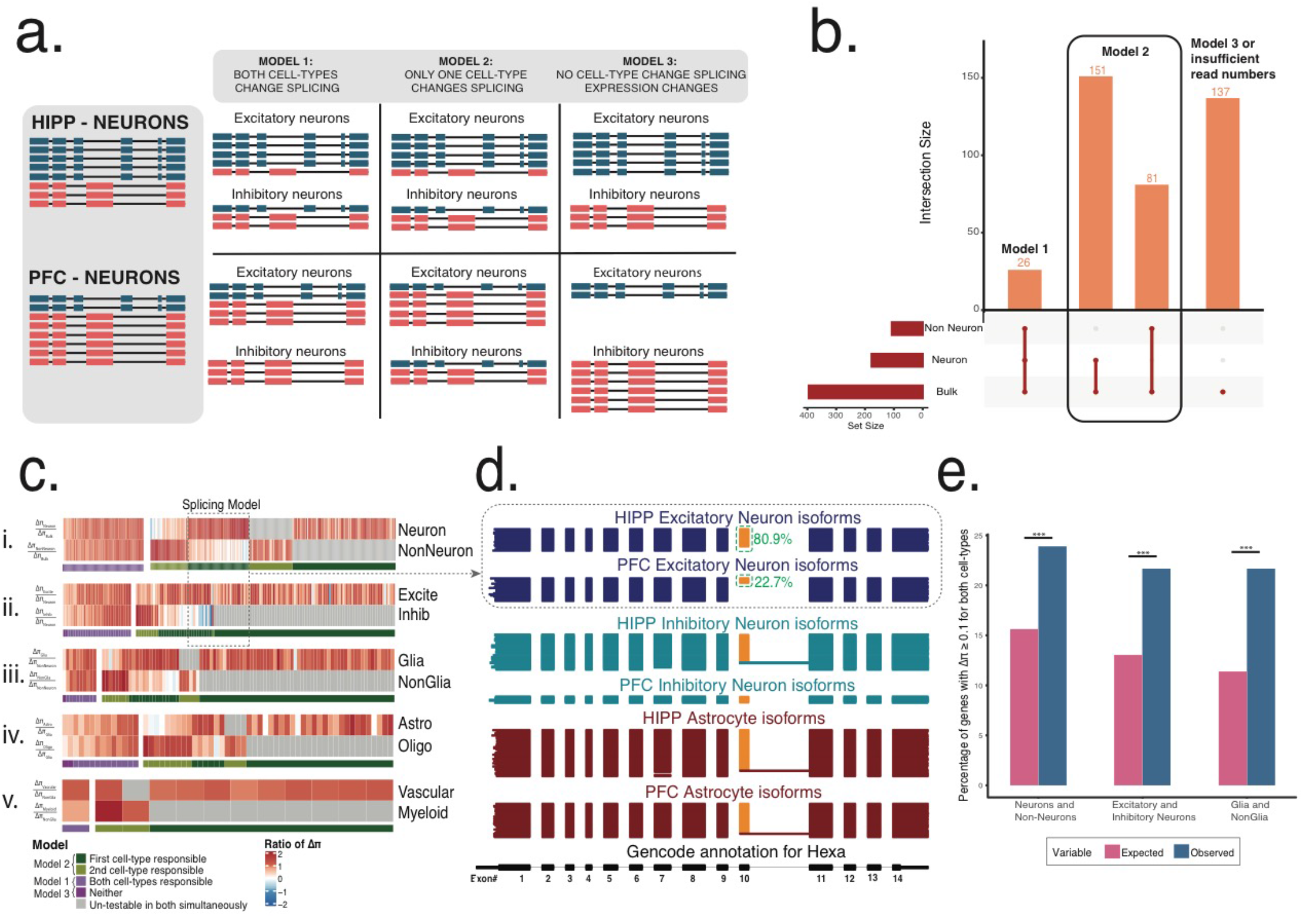
Three models of alternative splicing followed by differentially spliced genes across brain regions. **a.** Schematic of three models that explain splicing changes between any two categories (e.g. HIPP neurons versus PFC neurons) **b.** Upset plot of DIE genes in bulk, neuron, and non-neuron. **c.** Five gene×celltype heatmaps clustered by the ratio of Δπ of an individual cell-subtype to a parent cell-type. Each vertical line indicates the ratio of Δπ for a single gene. Grey lines indicate lack of sufficient depth or lack of expression. Clusters of genes are colored by whether both cell-types show similar relative Δπ to the parent (purple, Model I, Model III) or whether one cell-type explains most of the splicing changes (Model II). **d.** Hexa gene representing an example of Model II. Each horizontal line in the plot represents a single transcript colored according to the cell-type it is represented in. Therefore, blocks represent exons and whitespace represents intronic space (not drawn to scale). Orange exon represents alternative inclusion **e.** Barplots indicating percent of genes with |Δπ| >= 0.1 for two concurrently assessed cell-types. Pink bar indicates expected levels assuming independence, while blue bars represent observed levels.

We analyzed neuronal and non-neuronal reads separately and cross-referenced the results of the 395 genes with DIE. 26 genes (6.6%, FDR <= 0.05) had DIE in neurons and non-neurons under the Both-Cell-Types-Model, 151 (38.2%) in neurons only and 81 (20.5%) in non-neurons only under a Single-Cell-Type-Model, and 137 (34.7%) that were either too low in expression for testing (Methods) in both neuronal and non-neuronal cells or followed the No-Cell-Type-Model (Fig 3b). To distinguish these two explanations for the 137 genes, we considered genes with ΔΠ≥0.1 irrespective of p-values for the 395 genes. Specifically, we calculated the ratio of ΔΠ in a finer subtype to the ΔΠ in the composite cell type. After dividing all cells (composite) into neurons and non-neurons (finer level), 75% (± 2.3, SEp) of bulk DIE events were traced to only neurons or only non-neurons (Single-Cell-Type-Model, green) and 24.4% to both (Both-Cell-Types-Model, light purple, Fig 3ci). A single gene followed the No-Cell-Type-Model (0.3%, dark purple). Similarly, when dividing neurons into excitatory and inhibitory subtypes, we found 78.8% (±2.97) for the Single-Cell-Type-Model, 19.58% for Both-Cell-Types-Model and 1% for No-Cell-Type-Model (Fig 3cii). When separating composite non-neuronal cells into glia and non-glia, and again when separating glia into astrocytes and oligodendrocytes, and then non-glia into vascular and myeloid cells, we observe similar trends. The Single-Cell-Type-Model was more prevalent than the Both-Cell-Types and No-Cell-Type Models (Fig 3c). These observations were replicated in replicate 2, despite differences in cell-type proportions between replicates (Fig S7a-c). In summary, the No-Cell-Type-Model is rare, representing 0.3-3.28% for the different cell group divisions. Extending this observation to the 137 genes above (Fig 3b), one gene (0.3% of 395) likely represents the No-Cell-Type-Model whereas the other 136 can be attributed to low read depth.

An example of the dominant Single-Cell-Type-Model is Hexosaminidase A (*Hexa*), which encodes a lysosomal enzyme subunit and is implicated in Tay-Sach’s disease, a neurodegenerative disorder lethal in infancy^42^. Previously, only one annotated isoform existed for this gene. However, PFC excitatory neurons show significantly diminished inclusion of an internal exon (from 81% inclusion to 22%), thus expressing a novel isoform, while other cell types show no difference between HIPP and PFC. Manual validation classified this novel isoform as NMD, indicating brain-region and cell-type specific NMD (Fig 3d).

### Brain regions can override cell-type specificity for a subset of genes, possibly through microenvironmental influence

Despite the prevalence of the Single-Cell-Type-Model, the Both-Cell-Types-Model is still common. To avoid circular reasoning, we considered all genes sufficiently expressed (Methods) in neurons and non-neurons in both brain regions. Concurrent regional DIE differences (ΔΠ≥0.1) in both neurons and nonneurons occur more often than expected by chance 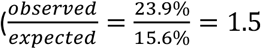, *p* ≤ 2.2*e*^-16^, Fisher’s two-sided exact test). This trend is also conserved in excitatory and inhibitory neurons (*p* ≤ 2.2*e*^-16^) as well as in non-neuronal glia and non-neuronal non-glia (Fig 3e). Two non-mutually exclusive models may underlie this observation – microenvironment and cell origin. Firstly, HIPP and PFC interneurons originate in the ganglionic eminences^31,33^, while excitatory neurons do not. Thus, splicing similarities between HIPP EN and IN that are different from PFC EN and IN might be imposed by the regional microenvironment. Secondly, considering neurons and glial non-neurons, their common descendance from radial glial stem cells may underlie cases of brain-region specific regulation that overrides cell-type specificity.

### Cell types endogenous to one brain region have distinct splicing signatures

We traced the contribution of individual cell types to bulk DIE. Region-specific DIE was clearly traceable 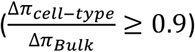 in 73.4% (n=395) of the cases, while 10.4% had 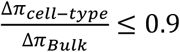 in all cell types (Fig S7d-e). The remaining 16.2% had low read counts in all cell types. Some genes truly have a regional component i.e. concurrent changes across multiple cell types (Fig S7e). The low depth and ΔΠ ratio could arise from cell types only present in HIPP but not PFC. Indeed, reads originating from granuleNB, RGL, ependymal, and CPE clusters have a higher ratio in the 26.4% of genes for which DIE was not explained (grey) than in the genes where DIE was explained (yellow) (Fig S7f). For example, for the *Fxyd1* gene, CPE cells in HIPP had a different splicing signature from astrocytes in HIPP and PFC, leading to regional DIE (Fig S7g). These observations warrant further exploration of each cell type’s splicing signatures.

### Choroid plexus epithelial cells (CPEs) generate distinct isoforms predominantly through alternative TSS

We performed DIE tests in pairwise comparisons of HIPP cell types. DIE was most frequent for neuron vs. non-neuron comparisons in HIPP (Fig 4a), and this was confirmed in PFC (Fig S8). High percentages were also seen in some comparisons between non-neuronal cell types. Interestingly, comparisons between non-neuronal cell types showed higher DIE than those observed within neurons (Fig 4b). Importantly, nonneuronal comparisons involving CPEs clearly had the highest DIE fractions. CPEs are cerebrospinal fluid secreting ependymal cells in cerebral ventricles, and alternative splicing in CPEs relates to disease^9,43^ (Fig 4c). Surprisingly, TSS choice (compared to exons and polyA-sites) largely explained the isoform regulation of CPE cells (Fig 4d). Furthermore, CPE-associated transcripts strongly favored an upstream TSS compared to the non-CPE transcripts (70/93 genes, Bernoulli p=3×10^-7^). This can allow for CPE-specific post-transcriptional modifications^44^, translation initiation, and transcription factor control of gene expression (Fig 4e).

**Figure 4 -.**
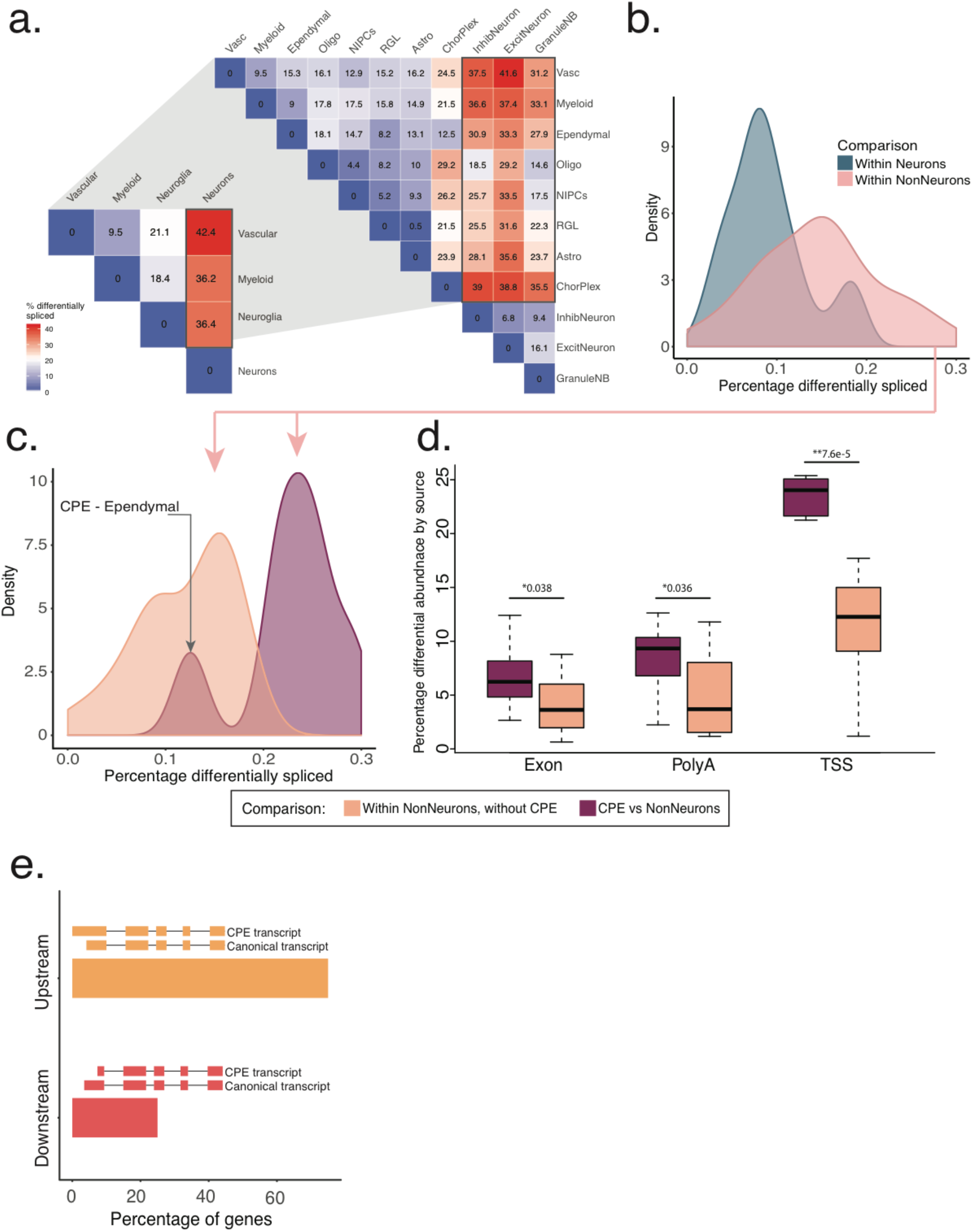
Choroid plexus epithelial cells (CPEs) generate distinct isoforms predominantly through alternative TSS. **a.** Triangular heatmap representing percentage of significant DIE in pairwise comparisons at two levels of granularity. At the broad level: pairwise comparisons of neurons, non-neurons, myeloid cells and vascular cells. Zooming in, at the narrow level: neuronal and non-neuronal categories are broken up into their constitutive cell-subtypes **b.** Density plot of percentage of significant DIE in pairwise comparisons from 4A, broken down by two categories: within neurons and within non-neurons. **c** Density plot showing DIE within non-neurons (pink region, fig 4b) broken up further by comparisons that either include (purple) or exclude (orange) choroid plexus epithelial (CPE) cells **d.** Boxplots of percentage significant genes in non-neuronal comparisons including and excluding the CPE, broken down by part of the transcript (TSS/splice-site/PolyA) responsible for splicing changes. X-axis indicates the type of test conducted **e.** Barplot showing the percentage of genes for which the CPE transcripts are either upstream or downstream of non-CPE transcripts

### Clustering on long-read data recapitulates short read cell-type assignments

We clustered hippocampal cells using their isoform expression similarities. Compared to 3’seq clustering, glial, myeloid and vascular clusters were similarly defined (Fig S9a-b, 4a). Jaccard similarity index analysis between short-read and long-read clusters showed high concordance for broad-level classification (Fig S9c). Additionally, isoforms of some genes are better resolved with long reads than with than 3’-expression short reads, including for distinguishing neurons and non-neurons (e.g. *Pkm, Clta, H3f3b*), or mature and immature neurons (e.g. *Cdc42, Srsf3, Thra*, Fig S9d). However, despite striking differences in isoform expression within neuronal subtypes, differences between isoform-derived clusters and shortread derived clusters remained. Long-read clustering successfully separated CA1 from CA3 neurons (i.e. short-read EN1 vs EN2) but did not separate all cells of the more immature IN2 cluster from mature granule neuroblasts i.e. GranuleNB-2 (Fig S9e-f). Such differences could be explained by cell subtype specificity in isoforms only, or reduced sequencing depth of isoforms.

### Relative isoform expression differences during development reflect dynamic changes in function

Using Slingshot^45^ on a subset of hippocampal cells, we recovered the radial-glia-like (RGL) to excitatory-neuron developmental lineage. From RGLs to NIPCs only 5.1% of tested genes showed DIE (n=73; 95% CI=[4.03,6.34]). However, threefold more did so from NIPCs to GranuleNB (n=359, 95% CI=[15.81,19.10]) and then from GranuleNB to excitatory neurons (n=423, 95% CI=[14.72,17.54]) (Fig 5a). Gene ontology (GO) analysis^46,47^ revealed isoform changes in the splicing machinery itself in earlier steps, i.e. from RGL to NIPCs (for *Luc7l2, Hnrnpa2b1, Snrnp70, Srsf2, Srsf5, Srsf6, Srsf7, Rbm3*; Figs 5b, S10-11). However, as granule neuroblasts matured to excitatory neurons, DIE was additionally associated with synapse formation and axon elongation (*Snap25, Snca, Syp, Dbn1, Cdc42, Nptn, Gap43*) among others (Figs 5c-d, S10-11).

**Figure 5 -.**
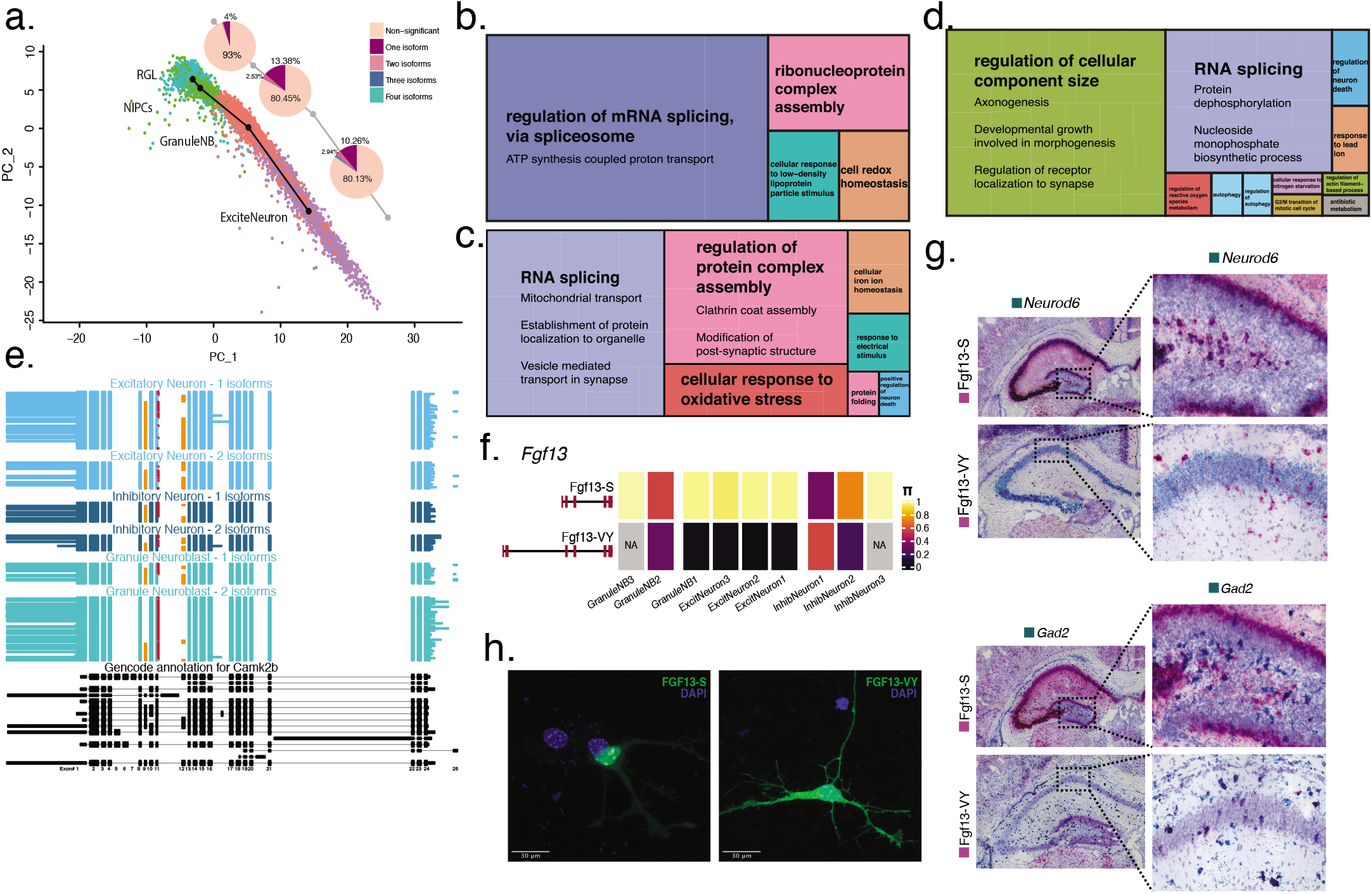
Relative isoform expression differences during development reflect dynamic changes in function. **a.** Splicing changes seen at every transition step of neuronal differentiation trajectory. Pie chart indicates the number of isoforms needed to be considered in order to reach the |Δπ| ≥ 0.1 threshold for a gene to be considered significantly DIE **b.** Treemap of condensed gene ontology (GO) terms for the transition from RGL to NIPCs in the neuronal differentiation trajectory, with size of boxes corresponding to number of significant terms associated with the GO category **c.** Treemap of condensed GO terms for the transition from NIPCs to GranuleNB (GNB1, GNB2, GNB3) **d.** Treemap of condensed GO terms for the transition from NIPCs to Excitatory Neurons (EN1, EN2, EN3) **e.** Hippocampal cell-type specific isoform expression in Camk2b gene. Each horizontal line in the plot represents a single transcript colored according to the cell-type it is represented in. Orange exons represent alternative inclusion, red extension of exon 11 represents a 3nt addition **f.** π value of the S and VY isoforms for Fgf13 gene across hippocampal neuronal cell types **g.** Basescope images of Fgf13-S and Fgf13-VY isoform expression in the hippocampus (pink stain), with simultaneous staining for excitatory neurons (Neurod6) and inhibitory neurons (Gad2). Zoomed in versions of the dentate gyrus for the S isoform, and CA1 region for the VY isoform. **h.** Subcellular localization of overexpressed Fgf13-S isoform in nucleolus and Fgf13-VY isoform in cytoplasm.

Importantly, many exon inclusion levels are altered in the transition from dentate gyrus (DG) granuleNB to more mature EN in CA1 and CA3. The synaptic Calcium/Calmodulin-dependent protein kinase II Beta (*Camk2b*) has enzymatic and structural roles in neuronal plasticity^48^. Embryonic *Camk2b* exploits a 3nt addition to exon 11 and exon-12 exclusion, translating an Alanine instead of Valine^49^. The embryonic form (red exon-11 extension, exon-12 exclusion, Fig 5e) dominates in more immature granuleNB2 compared to granuleNB1. However, this isoform persists infrequently in mature neuronal types (EN1, EN2, and IN1) indicating cell-type specificity during developmental regulation. Moreover, the additional 3 nucleotides (red in exon 11) co-occur with exon 12, which has not been reported. Also, the first alternative exon (exon 9) increases inclusion as cells differentiate. Furthermore, exon 9 coordination with exon 12 defines cellsubtype differences between EN and IN. All three splicing events occur in the actin binding domain of the CaMKIIß structure and encode several confirmed phosphorylation sites^50^. Thus, exon coordination among distinct cell types could indicate cell-type specific morphological changes in the actin cytoskeleton, for instance in spine dynamics^51^ (Fig 5e).

Discs Large Homolog Associated Protein (*Dlgap4*) encodes the post-synaptic protein SAPAP4, which is critical for functional neuronal network development, and *Dlgap4* mutations are linked to neuropsychiatric disorders^52^. We find that neuronal cells employ two exons, whereas non-neuronal cells predominantly express only one exon. As cellular identity changes from DG cell types (GranuleNB, IN3) to mature neuronal cell-types in the CA region (EN1, EN2, EN3, IN1, IN2), exon inclusion switches from a single exon to both (Fig S12a-b). Similarly, in another synaptic gene, Neuroplastin (*Nptn*), involved in longterm potentiation, neuroglia, vascular, myeloid, and immature DG cell types predominantly express a single 9nt micro-exon. Mature excitatory neurons and interneurons (EN1, EN2, EN3, IN1, IN2) conversely employ an upstream acceptor, adding four amino acids that code for the cytoplasmic domain, likely relevant for protein-protein interactions^53,54^ (Fig S12b).

### Hippocampus enriched developmental gene Fibroblast growth factor 13 (*Fgf13*) shows neuronal subtype specific TSS

*Fgf13* exhibited high DIE (ΔΠ > 0.5) across multiple neuronal cell-type and subtype comparisons. *Fgf13* reaches peak expression, specifically in HIPP, at our investigated time point (P7)^55^. *Fgf13* is a neuronal developmentally regulated gene and lethal when knocked-out^56,57^. It has multiple intracellular roles including regulation of voltage-gated sodium channels^58–60^, rRNA transcription^61^, and microtubule stabilization^55^. Of the various *Fgf13* isoforms^62^, two isoforms with distinct TSS dominate during brain development^55,63^. We find that *Fgf13* is particularly alternatively spliced between excitatory and inhibitory neurons (Fig 5f). The downstream-TSS isoform, Fgf13-S, is the major HIPP isoform across all excitatory types, and immature inhibitory types. Conversely, the upstream-TSS isoform, Fgf13-VY, is partially seen in DG neuroblasts and dominates in inhibitory interneuron classes. This was confirmed using Basescope analysis with *Neurod6* staining for excitatory neurons and *Gad2* staining for inhibitory neurons (Fig 5g, Methods). The isoforms also differ in subcellular localization: Reflecting its role in regulating protein synthesis^61^, Fgf13-S is primarily localized to the nucleolus, whereas Fgf13-VY is present throughout the cytoplasm, consistent with its known role in regulating voltage-gated sodium channels^59^ (Fig 5h).

### Synaptic genes associated with vesicle transport show splicing differences in developing hippocampal neurons

Many synaptic genes have low expression in developing HIPP. However, when sufficiently expressed (>50 reads), we observe splicing differences between neuronal subtypes. For example, Synaptosome associated protein-25 (*Snap25*) bridges synaptic vesicles to the plasma membrane during exocytosis^64^. Two mutually exclusive exons defined two developmentally regulated protein isoforms (Snap25-a and Snap25-b). Concordant with the literature^64–66^, we find expression of Snap25-b at P7 and find that CA3 excitatory neurons (EN3) have a higher proportion of Snap25-b transcripts than EN1 and EN2, which are localized in CA1 and subiculum. However, we see added cell-type specificity in exon inclusion as cells mature from granuleNBs to EN. GranuleNBs have higher abundance of Snap25-a whereas more mature excitatory neurons switched to Snap25-b. Interestingly, interneuron precursors and Cajal-Retzius cells (IN1,IN3) rely more on Snap25b than their excitatory counterparts, and thereby seem to rely more heavily on larger primed vesicle pools^67^ (Fig S13b). Alternatively, interneurons could switch from Snap25-a to Snap25-b before excitatory neuron development.

We also observe synaptic genes with key splicing differences between neurons and non-neurons. Clathrin light chain A and B (*Clta, Cltb*) encode synaptic proteins involved in clathrin-coated vesicle endocytosis, and work alongside *Epsin1* in cargo selection in endocytosis^68^. For all three genes, exon inclusion distinguishes neurons from non-neurons and granuleNB from more mature EN. For *Clta*, an additional exon distinguishes neuronal subtypes (Fig S14a-b). While the role of these *Clta* exons is unknown, the neuronal specific insertions in the clathrin light chain may play a role in the association with the slow axonal transport of clathrin^69^.

### Slide isoform sequencing (sliso-Seq) to delineate spatial localization of splicing changes

To ground our observations in a spatial sense, we generated 10X Genomics Visium spatial transcriptomics data from a P8 sagittal section. Alignment with single-cell short-read HIPP data confirmed the spatial localization of excitatory neurons in the CA regions and subiculum (Fig 6a), and alignment with the PFC data confirmed the excitatory precursors in distinct cortical layers (Fig 6b). We then devised a long-read sequencing approach for spatial transcriptomics (Methods). To validate regional specificity of isoform expression using orthogonal techniques, we correlated ΔΨ values between composite HIPP and composite PFC single-cell data with ΔΨ values of hippocampus vs. cortex using long-read sequencing. Based on single-cell data, we focused on 40 exons with region-specific splicing patterns and without alternative acceptors/donors (Methods). Overall, we found strong concordance: for 85%, both single-cell and spatial HIPP vs PFC splicing differences point in the same direction (Bernoulli probability <= 3.5e-06) (Fig 6c). Additionally, we confirmed neurodevelopmental exon inclusion switches in *Pkm* and *Clta*, where the non-neuronal and developmental exons from the single-cell data were enriched in the DG and in the choroid plexus of the spatial data (Fig S14, Fig 6c). For *Snap25*, the neurodevelopmental switch from Snap25-a to Snap25-b in single-cell data (compare Fig S13b), occurs in a posterior-to-anterior gradient in spatially mapped exons (Fig 6d). This supports the idea that the microenvironment can dictate brainregion specific splicing for some genes. Also, the hypothesis that interneurons selectively switch isoforms before excitatory neurons seems unlikely.

**Figure 6 -.**
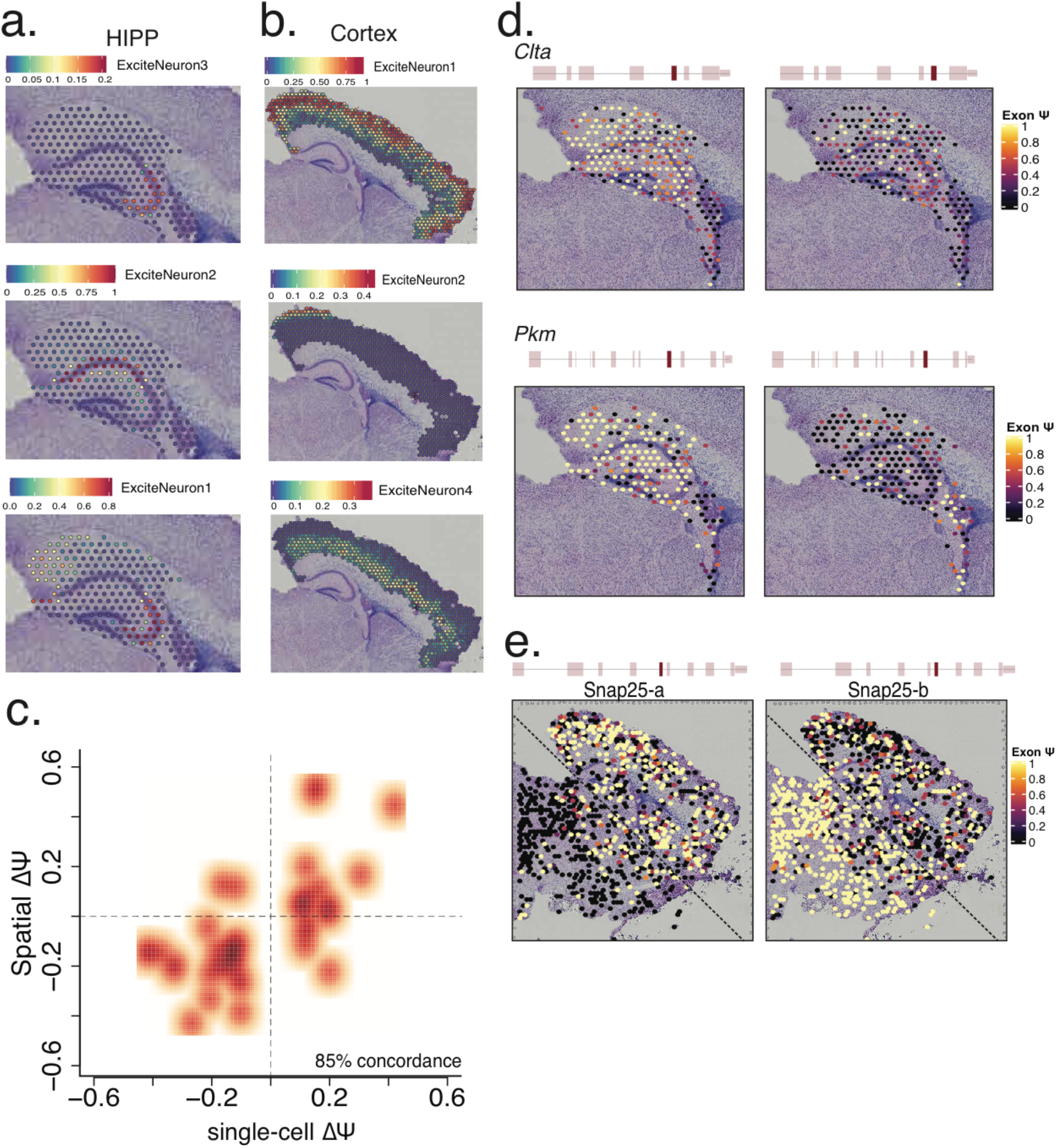
Slide isoform sequencing (sl-ISO-Seq) confirms spatial localization of splicing changes. **a.** Spatial localization of hippocampal single-cell excitatory neuron subtypes using gene-expression similarities to HIPP short reads. Color of spot indicates percentage of transcripts corresponding to indicated cell type **b.** Spatial localization of cortical excitatory neuron subtypes using gene-expression similarities to PFC short reads. Color of spot indicates percentage of transcripts corresponding to indicated cell type **c.** Scatter plot of the Δψ between HIPP and PFC from the single-cell data, and the Δψ between HIPP and cortex from 10X Visium spatial data (r^2^=0.6) **d.** Spatial distribution of alternatively spliced exons in *Clta* and *Pkm* genes in the hippocampal and choroid plexus regions. Color of spot indicates ψ values for each exon. **e.** Spatial distribution of two mutually exclusive, alternatively spliced exons in *Snap25* gene across the whole slide. Color of spot indicates ψ values for each exon.

## Discussion

Temporal and anatomic differences in alternative splicing are implicated in developmental changes in molecular function^70^. Our single-cell isoform data enable, for the first time, the illumination of alternative splicing regulation across cell types. We endeavored to generate cell-type, brain-region, and age-specific maps of AS to understand functional consequences of differential isoform expression (DIE). Our analysis hinges on a single-per-gene isoform test that considers a gene’s entire isoform repertoire and outperforms exon-based tests. Region-specific isoform testing revealed 395 genes with DIE between hippocampus and PFC, partially caused by isoforms novel to GENCODE. Manual validation by the GENCODE team using rigorous metrics lends credibility to this novel isoform detection. Thus, filtering these isoforms using bulk short reads, CAGE and polyA-site data provides a mechanism to automate genome annotation for isoforms.

We defined cell types underlying DIE between brain regions. In most cases and levels of granularity, we identified a finer cell type explanation of DIE, including the altered part of the transcripts. Despite inhibitory interneurons in HIPP and PFC migrating in from a common origin^31^, we observe a signature of coordinated DIE in excitatory and inhibitory neurons for a gene subset. Such gene subsets with coordination across cell types were observed at all investigated levels. Microexons are linked to neuronal cell types for their regulatory function^1,41^, however, in the case of *Nsfl1c* we find that their expression is not limited to neurons but instead exhibits regional regulation by being expressed only in HIPP. Thus, brain region can override cell-type specificity for a subset of genes, which may explain region-specific sQTLs^21^. The theoretical possibility that brain-region DIE in bulk arises purely from cell-type abundance or transcriptional activity differences is rarely observed. However, to which extent these observations persist in case-control studies of disease, across adult brain regions, and across species requires further studies.

Our results indicate that understanding the cell-type basis of sample-specific DIE requires a thorough understanding of cell-type specific DIE within each sample. This further warrants a brain-wide map of isoform expression at a single-cell level. Within the brain, isoform diversity in non-neuronal cell types has attracted less attention than in their neuronal counterparts. However, we find that non-neuronal cell types exhibit high pairwise DIE. Choroid plexus epithelial cells show particularly high differences from other non-neuronal (but also neuronal) cell types. Surprisingly this observation is largely caused by CPE-specific choices of predominantly upstream TSSs. This raises the question whether other highly specialized cell types in other brain areas exhibit similar complex alternative transcriptome mechanisms.

We notice the sheer functional diversity that AS lends to the splicing machinery, synaptic plasticity and vesicle mediated endocytosis. This motivates further investigation linking a spliceosomal-gene isoform to the splicing of its target gene, as well as the isoform state of synaptic genes in neuronal subtypes. The evidence for isoforms adding to cellular diversity is further bolstered by long-read clustering, which yields coherent cell-types albeit with discrepancies, which could be partially due to the sparsity of the isoform matrix due to lower long-read throughput. Alternatively, cell states could be defined by an isoform expression program that doesn’t correspond to 3’seq-based cell-type definition, highlighting the need for factoring in isoform expression to re-define traditional transcriptional cell-types.

Finally, we have devised the new method of Slide-isoform sequencing (sliso-Seq), which employs spatial transcriptomics and long-read sequencing. This allows anchoring the above results in a spatial view of the brain and reveals important biology such as the brain-wide coordination of the *Snap25* isoform switch. Looking forward, the integration of long-read single-cell and spatial technologies allows for the possibility of constructing 3-dimensional maps of isoform expression at single-cell resolution.

In summary, these results present an unprecedented view of cell-type specific full-length isoforms across brain regions, bringing us closer to a comprehensive isoform map of the brain. The software generated here is employable in a much larger setting and available as an <u>R-package</u>. It extends to case-control studies, broad sample comparisons, and spatially anchored comparisons beyond mouse or brain.

## Acknowledgments

We thank Jenny Xiang, Dong Xu, and Adrian Tan at the Genomics Resources Core facility at Weill Cornell Medicine for help with facilitating sequencing, and Ishaan Gupta for helping with initial processing of the samples.

## Methods

### Ethics statement

All experiments were conducted in accordance with relevant NIH guidelines and regulations, related to the Care and Use of Laboratory Animals tissue. Animal procedures were performed according to protocols approved by the Research Animal Resource Center at Weill Cornell College of Medicine.

### Animals and tissue isolation

C57BL/6NTac (n=6) female pups were quickly decapitated. For the single-cell experiments (P7; Rep1 n=1, Rep2 n=2), the brains were removed and placed on a stainless-steel brain matrix for mouse (coronal repeatable sections, 1mm spacing), and the prefrontal cortex and hippocampus were dissected^71^ in cold PBS solution (Fig S15a-f). Brain tissues from both hemispheres were pooled in one sample. After dissection, tissues were snap frozen in dry ice until processing. For the 10X Visium spatial experiment (P8; n=1) brains were fresh-frozen and embedded in OCT. For the Basescope (P7; n=2) experiments, the brains were transcardially perfused, immersion fixated and cryo-protected (15% and 30% Sucrose in phosphate-buffered saline) each over night before being embedded in OCT

### Tissue disassociation

Following recommendations from 10x Genomics (Cat#CG00055 Rev C) dissected hippocampus and prefrontal cortex tissue was placed into 2.5 ml Hibernate E/B27/GlutaMax medium (BrainBits cat#HEB) at Room Temp until all samples were dissected. HEB medium was removed and replaced with 2ml of 2 mg/ml activated papain (BrainBits cat#PAP) then incubated for 25 min at 37°C with gentle mixing. After allowing tissue to settle, papain was removed and replaced with 2mL fresh HEB medium and tissue was gently triturated 15–20 times using a wide-bore pipette tip and tissue left to settle. Supernatant was taken and filtered using a 30μm cell strainer (Miltenyi Biotec cat#130-041-407) into a collection tube. To the remaining tissue, another 2ml of fresh HEB medium was added and then triturated with a regular 1ml pipette tip an additional 10 – 15 times until tissue was completely disassociated. Supernatant was taken and filtered through a 30um cell strainer and added to the collection tube. Supernatant was then centrifuged at 400rcf for 2 min. The cell pellet was re-suspended in 1 – 3ml of neuronal culture medium NbActiv1 (BrainBits cat#Nbactiv1) depending on cell pellet size, filtered through a 30μm cell strainer (Miltenyi Biotec cat#130-041-407) and was subsequently diluted to 1,500 cells/μl in NbActiv1 for capture on the 10x Genomics Chromium controller.

### 10x Genomics single-cell capture

The disassociated cells were captured on the 10x Genomics Chromium controller according to the Chromium Single Cell 3’ Reagent Kits V2 User Guide (10x Genomics CG00052 Rev F) with the following modification. PCR cycles were increased, from the recommended ten cycles for recovery of 8,000 cells, to 16 cycles to target a yield of cDNA enabling simultaneous Illumina and PacBio library preparation.

### Illumina and Pacific Biosystems library preparation

Illumina library preparation was performed using 100 ng of amplified cDNA following the Chromium Single Cell 3’ Reagent Kits V2 User Guide (10x Genomics CG00052 Rev F) reducing final indexing PCR cycles to ten cycles from the recommended 14 cycles to increase library complexity. Sequencing for Replicate 1 was performed on HiSeq4000 according to 10x Genomics run mode, for Replicate 2, sequencing was performed on a NovaSeq S1 flowcell also following 10x Genomics run mode and Bulk RNA-Seq was performed on the Illumina NextSeq 500 with a 150 PE run mode. PacBio library preparation was performed with 500 ng of amplified cDNA using SMRTbell Express Template Prep Kit V2.0 (PacBio cat#PN: 100-938-900) to obtain Sequel II compatible library complex and was sequenced on a total 24 Sequel I SMRTcells with a run time of 10 hours and 20 Sequel II SMRTcells with a run time of 30 hours across samples and replicates.

### Modification of 10x Visium Illumina library preparation

Illumina compatible libraries were made from Visium derived cDNA using a modified protocol derived from NEB Ultra II DNA FS kit (NEB #E6177). Visium derived spatial cDNA was diluted to 100ng in 26μl and combined with 7μl NEBNext Ultra II FS Reaction Buffer and 2μl NEBNext Ultra II FS Enzyme Mix and incubated at 37°C for 15min, 65°C for 30min to obtain fragmented, end-repaired and A-Tailed cDNA. Samples was then subjected to double-sided size selection by following the Beckman Coulter SpriSelect (Cat# B23318) protocol with an initial ratio of 0.6x SpriBeads. Supernatant was then taken and additional SpriBeads added for a final ratio of 0.8x eluting to 35μl EB Buffer. Adaptor ligation using 10x Genomics protocol was performed by combining 2.5μl 10x Adaptor Mix with 30μl NEBNext Ultra II Ligation Master Mix and 1μl of NEBNext Ligation Enhancer with the previously end-repaired cDNA and incubated at 20°C for 15min. Single sided SpriBead cleanup was performed using a 0.8x ratio and eluted in 15μl EB Buffer. Finally cDNA library was amplified by combining adaptor ligated cDNA with 5μl 10x Genomics i7 Barcoded primer and 5μl of 1:5 diluted 10x Genomics SI Primer and 25μl NEBNext Ultra II Q5 Master Mix and cycled with the following thermocyler profile: 98°C 30sec, 12 cycles of 98°C 20sec, 54°C 30sec, 72°C 30sec, then final extension of 65°C for 5min, 10°C hold. Amplified library was again subjected to a double-sided size selection using an initial ratio of 0.6x SpriBeads, supernatant was then taken and additional SpriBeads added for a final ratio of 0.8x eluting to 35μl EB Buffer. Illumina Libraries were then checked for quality and sequenced on an Illumina NextSeq500 instrument according to guidelines.

### Size selection of Visium cDNA using blue pippin for Oxford Nanopore

Visium derived cDNA was size-selected using Sage Sciences Blue Pipin instrument to obtain cDNA fragments 800bp to 6000bp for efficient sequencing on Oxford Nanopore PromethIon Instrument. Samples were diluted to 1000ng in 30μl of TE buffer and combined with 10μl of Sage Loading Solution before loading into one lane of a 0.75% Agarose Blue Pippin Cassette (Cat# BLF7510). Samples were then separated according to protocol for a target range of 800bp to 6000bp, and target elution retrieved after 12 hours. Samples were then cleaned by Beckman Coulter SpriSelect Beads (Cat# B23318) using a 0.8x ratio and eluted 50μl Nuclease Free Water. Size-distribution was checked using Agilent Fragment Analyzer Large Fragment Kit (Cat# DNF-464-0500).

### PromethION library preparation and sequencing of Visium cDNA

Oxford Nanopore compatible library was produced using 350ng of either Sage Blue Pippin size-selected cDNA or non-selected cDNA derived from 10x Genomics Visium following the Genomic DNA by Ligation protocol (SQK-LSK109) from Oxford Nanopore with the following modifications. End-Repair was carried out omitting NEBNext FFPE DNA Repair and incubations extended to 10min at 20°C and 10min at 65°C. Loading inputs on the PromethION was increased to 150fmol and sequenced for 20 hours.

### Total hippocampus and PFC short-read Illumina library preparation

Illumina compatible libraries were produced from 1250ng total RNA using NEBNext Ultra II RNA Library Prep Kit (NEB Cat#E7770S) following manufactures protocol with the following modifications. Target insert size was 450 bp for compatibility with paired end 150bp sequencing mode. Number of PCR cycles was reduced to 6 to limit the effect of PCR aberrations on the final library. Sequencing was performed on the Illumina NextSeq 500 instrument.

### Generation of circular consensus reads

Using the default SMRT-Link parameters, we performed circular consensus sequencing (CCS) as follows with the following modified parameters: maximum subread length 14,000bp, minimum subread length 10bp and minimum number of passes 3.

### Primary Hippocampal Culture and Transfection

Primary dissociated hippocampal cultures were prepared as previously described^59^, with minor modifications. Briefly, the hippocampus from P0 mouse pups was dissected on ice, digested with 0.25% trypsin for 30 min at 37°C with DMEM (Sigma), and dissociated into single cells by gentle trituration. The cells were seeded at a density of 2.5–3.0 × 10^5^ cells per coverslip in Neurobasal-A (Sigma) supplemented with 10% (vol/vol) heat-inactivated FBS onto coverslips previously coated with 50 μg/mL poly-D-lysine (Sigma) overnight at 4°C and 25 μg/mL laminin (Sigma) for 2 h at 37°C. The cells were maintained in a humidified incubator in 5% CO2 at 37°C. After 24 h, the medium was replaced with Neurobasal-A supplemented with 2% B27 (Invitrogen), 1% FBS, 25 μM uridine, and 70 μM 5-fluorodeoxyuridine. After 6 days of in vitro (DIV) culture, the neurons were transiently transfected with 0.2 μg of plasmid DNA per coverslip using calcium phosphate. One day after transfection cultured hippocampal cells were fixed for 30 min with 4% paraformaldehyde, washed three times with PBS, and incubated for 5 min in DAPI solution. Imaging was performed with a Zeiss LSM 880 Laser Scanning Confocal Microscope using an oil immersion 63× objective. All images were collected at a 2,048 × 2,048-pixel resolution. EGFP fusion constructs were generated as previously described^59^.

### Alignment of bulk short-read data for junction validation

Illumina short reads for HIPP and PFC were aligned to the reference genome (mm10) using STAR using the following parameters:

--outFilterMultimapNmax 1 --outFilterIntronMotifs RemoveNoncanonical --outFilterMismatchNmax 5 -- alignSJDBoverhangMin 6 --alignSJoverhangMin 6 --outFilterType BySJout --alignIntronMin 25 -- alignIntronMax 1000000 --outSAMstrandField intronMotif --outSAMunmapped Within --runThreadN 32 --outStd SAM --alignMatesGapMax 1000000

### Alignment of single-cell short read data and analysis

The 10x cellranger pipeline (version 3.0.0) was run on the raw Illumina sequencing data to obtain single-cell expression matrices. For replicate 1, the raw expression matrices obtained through cellranger were used along with the DropletUtils package^72^ to acquire ‘eligible’ barcoded single cells (FDR <= 0.001) with UMI counts that fell below cellranger’s filtering cutoff. These barcodes were incorporated into new matrices for importing into Seurat (v3.1). For both hippocampal replicates and the first PFC replicate, cells that had unique gene counts over 5,000 or less than 700, and greater than 20% mitochondrial gene expression were removed from further analysis. To adjust for the lower mean reads/cell for the second PFC replicate, the cutoff for minimum number of genes per cell was lowered to 350. Filtering on these parameters yielded 14,433 single cells for the hippocampus across two replicates, and 10,944 single cells for the PFC. We then used Seurat’s “merge” feature^73^ to combine the replicates for each brain region. The number of UMIs, percentage of mitochondrial gene expression were regressed from each cell and then the gene expression matrix was log normalized and scaled to 10,000 reads per cell. Next, we clustered all the cells using 30 principal components (PCs) using the Louvain algorithm with a 0.6 resolution.

### Alignment of spatial short read data and analysis

The 10X spaceranger pipeline was run on raw Illumina sequencing data to obtain spatial expression matrices. Seurat’s spatial analysis functions were used to obtain gene expression similarity clusters and identify barcodes corresponding to various brain regions.

### Integrated analysis with published data to identify cell-types

Published RNASeq P30 mouse brain data from Allen Brain Atlas^30^ was used as a reference to identify cell identities of clusters based on shared gene expression patterns. Since the Allen institute data was generated using the SmartSeq2 protocol, Seurat’s integrated anchor feature^27^ was used to align the two datasets and transfer cell-type labels (Fig S1e-f, S2e-f).

### Integrated analysis with spatial transcriptomics data to identify cell-types

P7 HIPP single-cell data was used as a reference to transfer labels onto P8 spatial transcriptomics data in the barcoded region corresponding to the hippocampus using Seurat’s integrated anchor feature^27^ using default parameters.

### Single-cell trajectory analysis

The velocyto python package^74^ was used to obtain .loom files from both replicates of HIPP and PFC singlecell data. After importing the UMAP co-ordinates of the datasets, the scVelo^75^ package and tutorial with default parameters were followed to acquire velocity plots (Fig S1c-d, S2c-d). The cells involved in neurogenesis and neuronal differentiation in the dentate gyrus and hippocampus were subsetted based on cell-type identity, and slingshot trajectory analysis^76^ was conducted on its first two principal components in an unsupervised manner (Fig 5a).

### Alignment of Pacific Biosciences long read data

Long read CCS fastqs sequences with PacBio were mapped and aligned to the reference genome (mm10) using STARlong and the following parameters:

--readFilesCommand zcat --runMode alignReads --outSAMattributes NH HI NM MD --readNameSeparator space --outFilterMultimapScoreRange 1 --outFilterMismatchNmax 2000 --scoreGapNoncan −20 -- scoreGapGCAG −4 --scoreGapATAC −8 --scoreDelOpen −1 --scoreDelBase −1 --scoreInsOpen −1 -- scoreInsBase −1 --alignEndsType Local --seedSearchStartLmax 50 --seedPerReadNmax 100000 -- seedPerWindowNmax 1000 --alignTranscriptsPerReadNmax 100000 --alignTranscriptsPerWindowNmax 10000

### Alignment of spatial transcriptomics data sequenced using Oxford Nanopore

Long reads sequenced on the ONT PromethION were mapped and aligned using minimap2 using the following parameters: -t 20 -ax splice --secondary=no

### Filtering of long reads for full-length, spliced, barcoded reads

This was done as described in our previous publication^26^ with added filtering in place for incomplete reads using published CAGE peaks and polyA site data^77,78^.

### Exon count assignment per cell-type

High confidence mapped and aligned reads were processed sequentially and compared to the Gencode M21 gene annotation using an in-house <u>script</u>. A Gencode-annotated exon (Fig S16a) was considered as being included in the read if both splice junctions were detected by the alignment. Due to the high error rate of ONT sequencing data, a variation of 3 bp was allowed for each splice junction, whereas for PacBio variation allowed was 2 bp, and overlapping exons were flagged (Fig S16b). In cases where a read spanned an exon, but its splice junctions were not detected, the exon was considered excluded (Fig S16c). Although ONT reads are known to often represent truncated versions of transcripts, terminal exons were counted but were only considered included when covered completely. Partially covered terminal exons were considered neither included nor excluded, and were discarded from the analysis (Fig S16d). Further, exon counts were aggregated to produce exclusion and inclusion rates for each particular cell group.

### Differential isoform tests

For isoform tests, a read was represented by a string denoting the TSS, introns, and polyA-site and each isoform was assigned an ID, with lower numbers corresponding to higher abundances. Next, counts for each isoform ID were assigned to individual cell-types. For each differential abundance test between two categories, genes were filtered out as ‘untestable’ if reads did not reach sufficient depth (25 reads/gene category). For genes with sufficient depth, a maximum of an 11 × 2 matrix of counts denoting *isoform* × *category* was constructed with the first ten rows corresponding up to the first ten isoforms, and the last row comprised of collapsed counts from all the other isoforms (if any). P-values from a χ^2^ test were reported per gene, along with a ΔΠ value per gene. The ΔΠ was constructed as the sum of change in percent isoform (Π) of the top two isoforms in either positive or negative direction. After these numbers were reported for all testable genes for a comparison, the Benjamini Hochberg (BH)^79^ correction for multiple testing with a false discovery rate of 5% was applied to return a corrected p-value. If this FDR p-value was <= 0.05 and the ΔΠ was more than 0.1, i.e. the change in percent inclusion of one or two isoforms was more than 10% between the two categories, then the gene was considered to be significantly differentially spliced.

### Differential TSS tests

For each read, we determined the TSS as described above. The counts of each TSS in two conditions (e.g. excitatory and inhibitory neurons) were then summarized in a *n* × 2 table of counts. As with the isoform tests, this table was reduced to a 11 × 2 table where the first ten rows represented the most abundant TSS and the 11^th^ represented all the other TSS. Testing was performed as described above for isoforms.

### Differential polyA-site test

For each read, we determined the polyA-site as described above. The procedure for testing was the same as described above for differential TSS.

### Differential exon tests

Testing for differential exon inclusion followed a separate framework. For testing differential inclusion between two categories, a 2 × 2 table of inclusion and exclusion counts per exon was constructed if it was not constitutively spliced (0.1 <= Ψ <= 0.9) without considering counts in individual categories. An exon was considered for testing if the expected counts followed the criteria^80,81^ 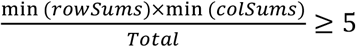, and the p-value from a χ^2^ test was reported, along with a ΔΨ value. The ΔΨ was constructed as the difference in percent spliced in (PSI/Ψ) across the two categories. After these numbers were reported for all testable exons for a comparison, since these tests are dependent, the Benjamini Yekutieli (BY)^36^ correction for multiple testing with a false discovery rate of 5% was applied to return a corrected p-value. If this p-value was <= 0.05 and the ΔΨ was more than 0.1 i.e. the change in Ψ was more than 10% between the two categories, then the exon is considered to be significantly differentially included.

### Isolating non-overlapping exons from single-cell data for correlation with spatial ONT data

To confirm region-specific DIE in a technologically orthogonal fashion, we used spatial data sequenced with Oxford Nanopore as opposed to single-cell data sequenced with PacBio. Because of the relatively high error rate, truncated transcripts (*200 bp difference in average length of ONT vs PacBio), and mis-mapping of overlapping splice sites, we chose to work with exons instead of full-length transcripts. We isolated exons from the single-cell PacBio data that showed region-specific DIE without p-value correction and extracted corresponding exons from the spatial data if they satisfied the following condition. If an exon was reported to have alternative donor and acceptor sites, it was only accepted if more than 90% of reads overlapping the exon mapped to that particular exon. We then used this list of exons to calculate region-specific DIE using spatial exon expression and correlated the ΔΨ values with the ΔΨ values from the single-cell data. Despite high technical variation, we got high concordance (85% change in the same direction, n=40)

### Building an enhanced annotation

We isolated polyadenylated, barcoded and spliced long-reads,

- whose TSS were within 50bps of a published CAGE-peak^77^
- whose mapped read-end fell within 50bps of a published polyA-site^78^
- whose intron-chains were inconsistent with any annotated transcript or any truncated version of an annotated transcript^82,83^
- whose internal exons (meaning both splice sites) were each supported by two or more single short reads (with >=2 splicing events) or paired-end Illumina read pairs from a bulk-sequencing experiment
- whose introns were each supported by two or more spliced Illumina reads from a bulk sequencing experiment
- who could not be interpreted as a truncated version of another such novel isoform

This resulted in an enhanced annotation, in which all added isoforms had a unique, previously absent intron-chain.

These isoforms as well as already annotated isoforms received read counts in each cell types according to the single-cell long-read dataset in each brain region. Novel isoforms intron-chains are by construction (see above) unique; however, annotated isoforms can differ only in their TSS or polyA-site. A long-read was therefore assigned to a GENCODE transcript that minimized the sum of abs(readEndMapping – annotatedIsoformEnd) and abs(readStartMapping – annotatedIsoformStart). In the case of a tie, only the divergence at the TSS was considered.

This annotation thus includes counts for thousands of isoforms in different cell types, such as “P7Hipp_OPCs” (representing oligodendrocyte precursor cells from a hippocampus at postnatal day 7) and “P7PFC_OPCs” (representing oligodendrocyte precursor cells from a pre-frontal cortex at postnatal day 7)

### Manual validation from the GENCODE team

Novel isoforms were manually verified and integrated into the GENCODE geneset^84^ in accordance with guidelines developed by the HAVANA group for the GENCODE / ENCODE projects^85^. Novel introns not previously identified within the GENCODE geneset were independently confirmed by intron predictions taken from the Ensembl RNAseq pipeline using ENCODE data^86,87^, whilst novel transcription start sites were validated based on the Cap Analysis of Gene Expression (CAGE) libraries generated by FANTOM^5^. Finally, each new transcript model was assigned a ‘biotype’ indicative of its presumed functional categorization, with the biotypes matching to those used in GENCODE as described by Frankish et al (2019)^84^.

### Validation of cell-type specific splicing using Basecope Duplex Detection Assay

Target transcriptomic regions for individual isoforms were isolated from the DIE analysis and probes were designed by Advanced Cell Diagnostics (ACD). Postnatal day 7 mice were transcardially perfused, and 12μm brain sections were processed according to the manufacturers recommendation for BaseScope Duplex Detection Assay (Protocol number 323800-USM, Advanced Cell Diagnostics, ACD) with a pre-treatment according to the protocol for fixed, frozen tissue (Protocol number 320534, Advanced Cell Diagnostics, ACD) with only 15 min of Protease Plus incubation. Images were taken using a Leica DM5500 B with a Leica DFC295 camera.

### Data and code availability

The source code for the methods and visualizations are available as functions in an R-package (https://github.com/noush-joglekar/scisorseqr) and processed single-cell and spatial data is available for download at www.isoformAtlas.com

