## Supplemental Material for "Cell-type, single-cell, and spatial signatures of brain-region specific splicing in postnatal development"

**Supplemental Figure 1**

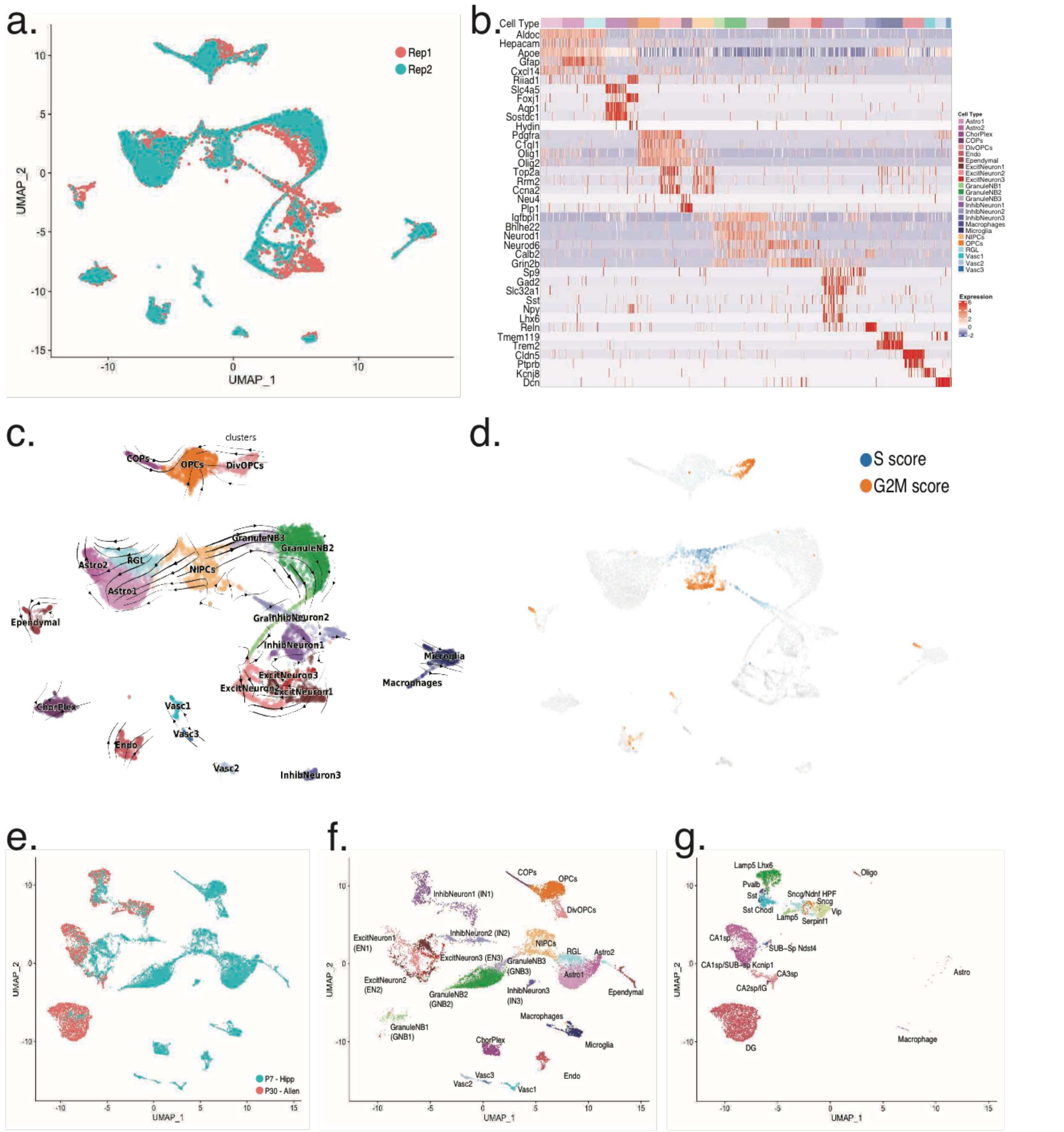


**S1. Identification of hippocampal cell-types based on single-cell short read gene expression** **a.** UMAP of hippocampal data colored by replicate **b.** Heatmap of marker gene expression across all identified cell types. Each horizontal band represents a gene, whereas each vertical line is a randomly chosen cell. Annotated blocks of cells at the top represent cell-types **c.** RNA velocity analysis of hippocampal single-cells **d.** Cell-cycle phase identified by RNA velocity analysis **e.** Integrated UMAP from aligning above single-cell P7 hippocampus data and P30 hippocampal data from the Allen Institute colored by time-point **f.** P7 hippocampal data extracted from S1E, colored by cell-type shown in S1B **g.** P30 hippocampal data extracted from S1E, colored by cell-types identified by the Allen Institute dataset.

**Supplemental Figure 2**

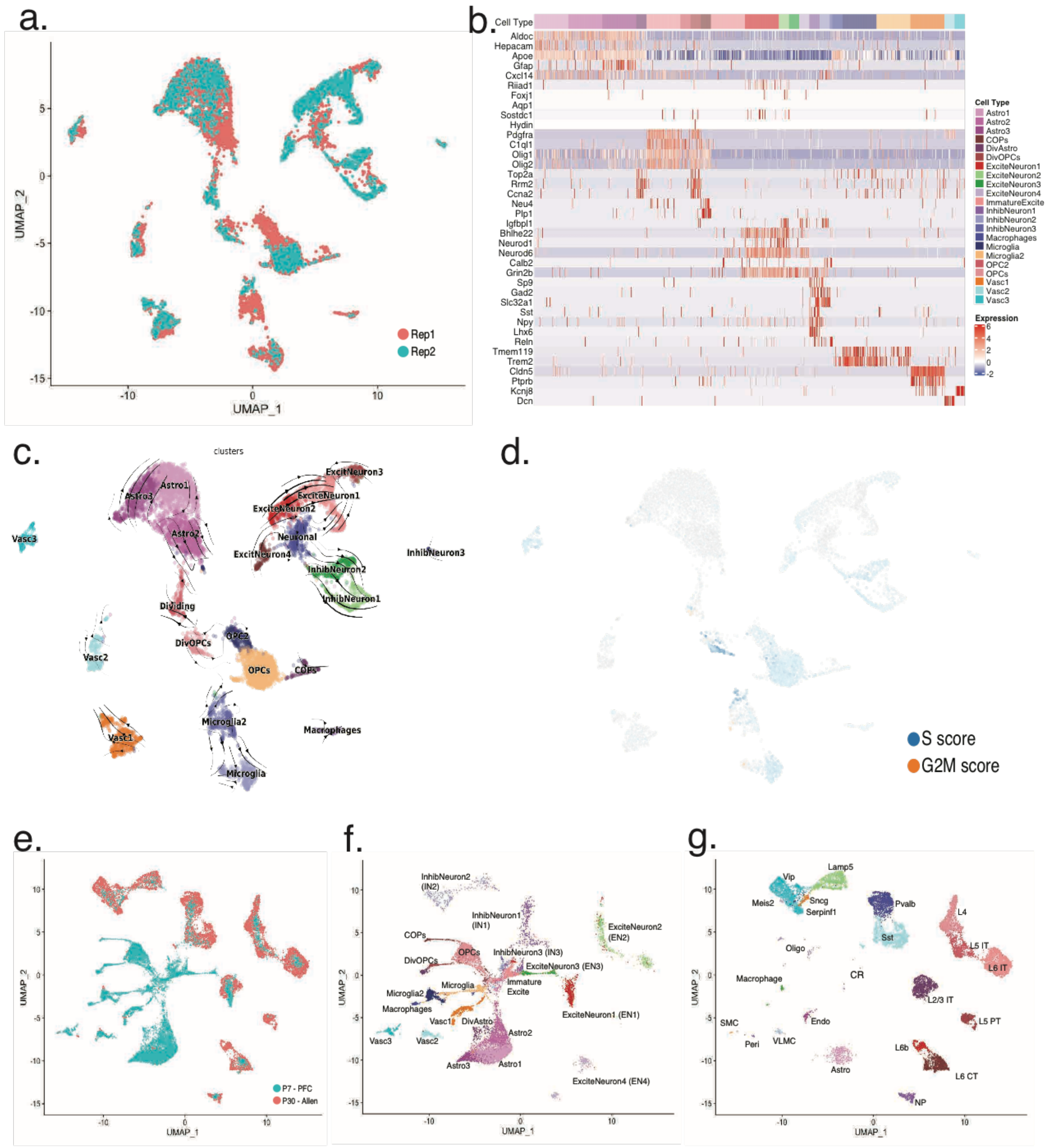


**S2. Identification of prefrontal cortex cell-types based on single-cell short read gene expression**

**a.** UMAP of hippocampal data colored by replicate **b.** Heatmap of marker gene expression across all identified cell types. Each horizontal band represents a gene, whereas each vertical line is a randomly chosen cell. Annotated blocks of cells at the top represent cell-types **c.** RNA velocity analysis of prefrontal cortex single-cells **d.** Cell-cycle phase identified by RNA velocity analysis **e.** Integrated UMAP from aligning above single-cell P7 PFC data and P30 cortex data from the Allen Institute colored by time-point. **f.** P7 PFC data extracted from S2E, colored by cell-type shown in S2B **g.** P30 cortex data extracted from S2E, colored by cell-types identified by the Allen Institute dataset.

### Supplemental Figure 3

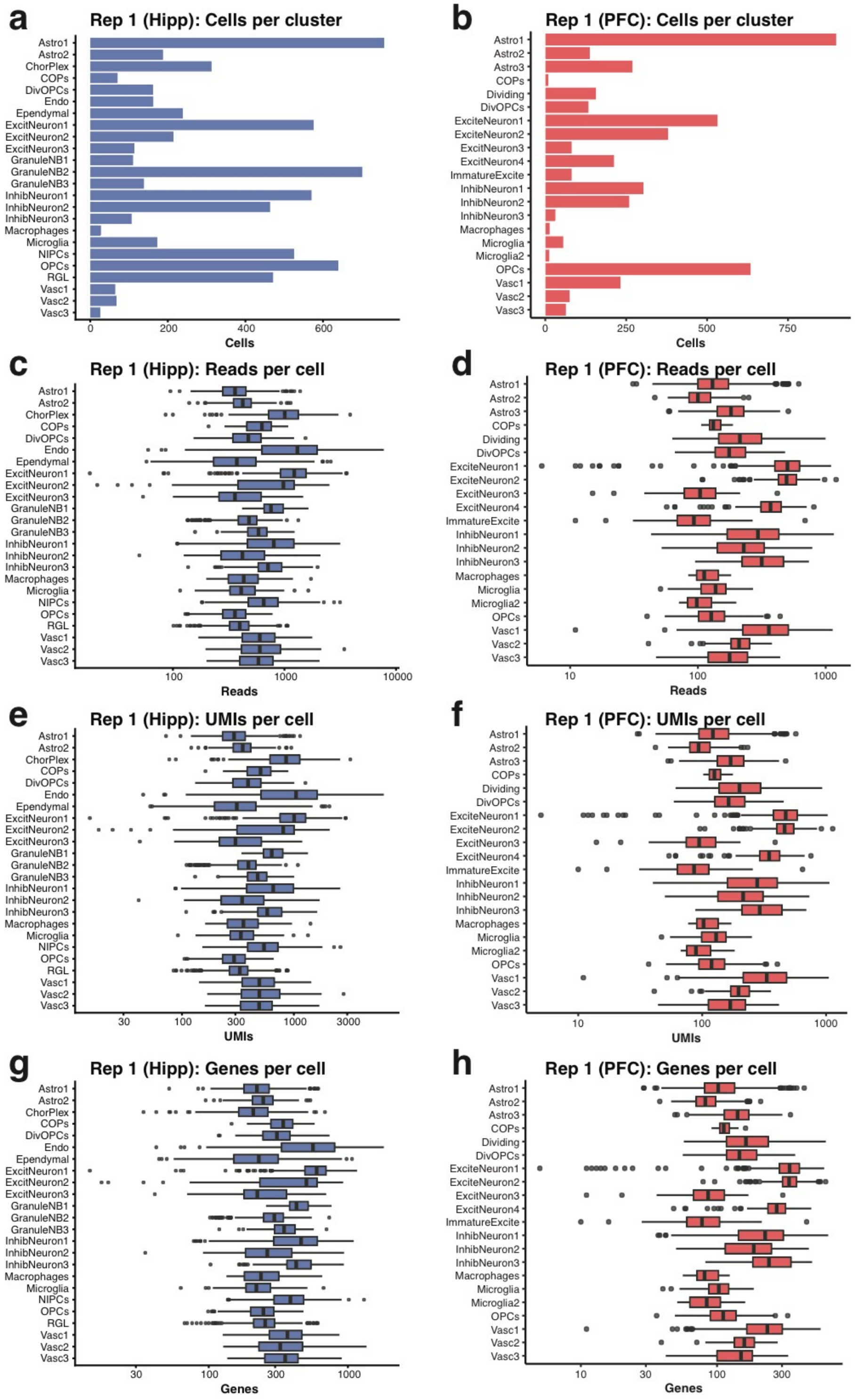


**S3. Long-read statistics for Replicate 1**

**a.** Barplot of cells per cluster identified in HIPP **b.** Barplot of cells per cluster identified in PFC **c.** Boxplot of reads per cell, grouped by cluster in HIPP **d.** Boxplot of reads per cell, grouped by cluster in PFC **e.** Boxplot of UMIs per cell, grouped by cluster in HIPP **f.** Boxplot of UMIs per cell, grouped by cluster in PFC **g.** Boxplot of genes per cell, grouped by cluster in HIPP **h.** Boxplot of genes per cell, grouped by cluster in PFC

**Supplemental Figure 4**

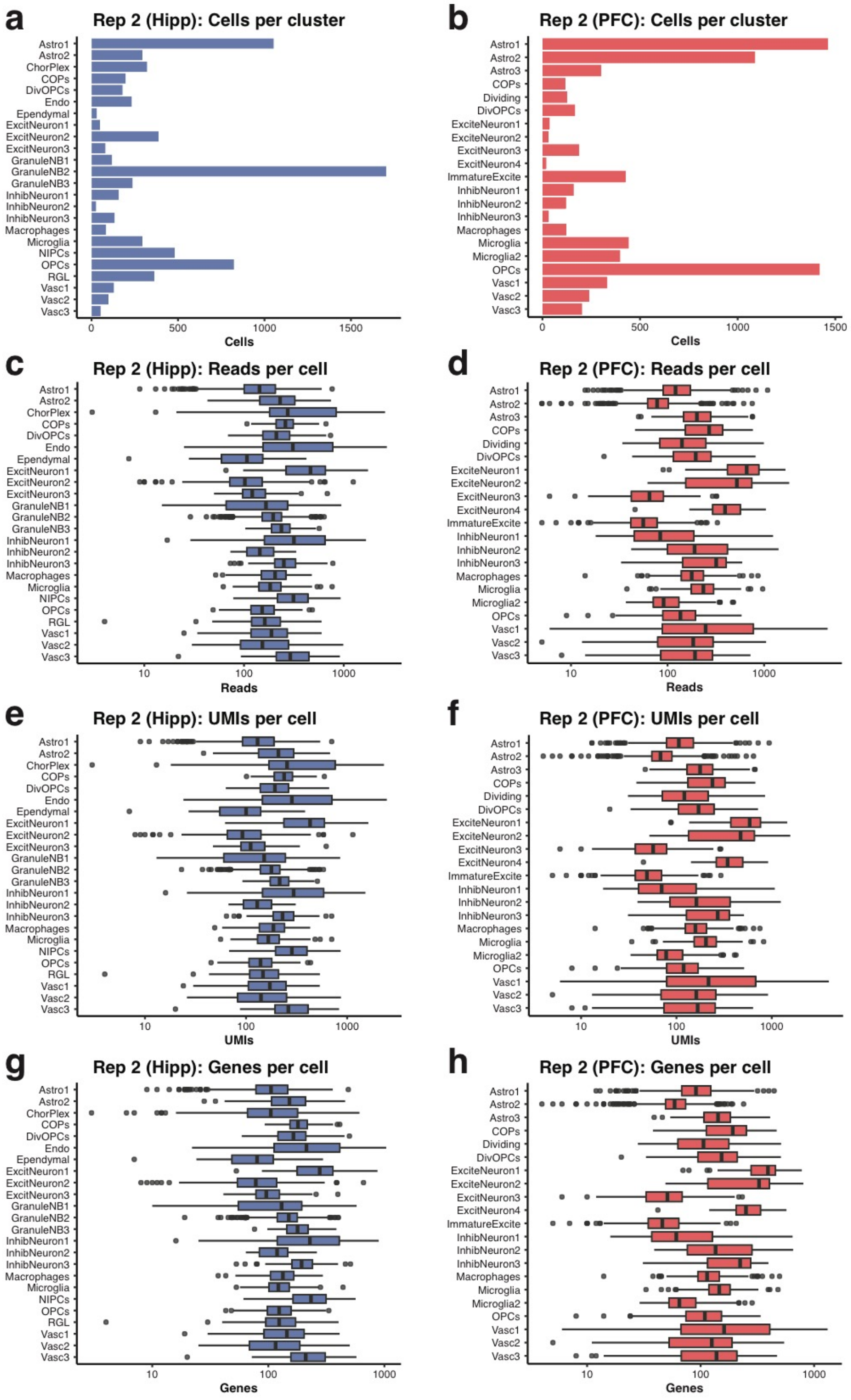


**S4. Long-read statistics for Replicate 2**

**a.** Barplot of cells per cluster identified in HIPP **b.** Barplot of cells per cluster identified in PFC **c.** Boxplot of reads per cell, grouped by cluster in HIPP **d.** Boxplot of reads per cell, grouped by cluster in PFC **e.** Boxplot of UMIs per cell, grouped by cluster in HIPP **f.** Boxplot of UMIs per cell, grouped by cluster in PFC **g.** Boxplot of genes per cell, grouped by cluster in HIPP **h.** Boxplot of genes per cell, grouped by cluster in PFC

**Supplemental Figure 5**

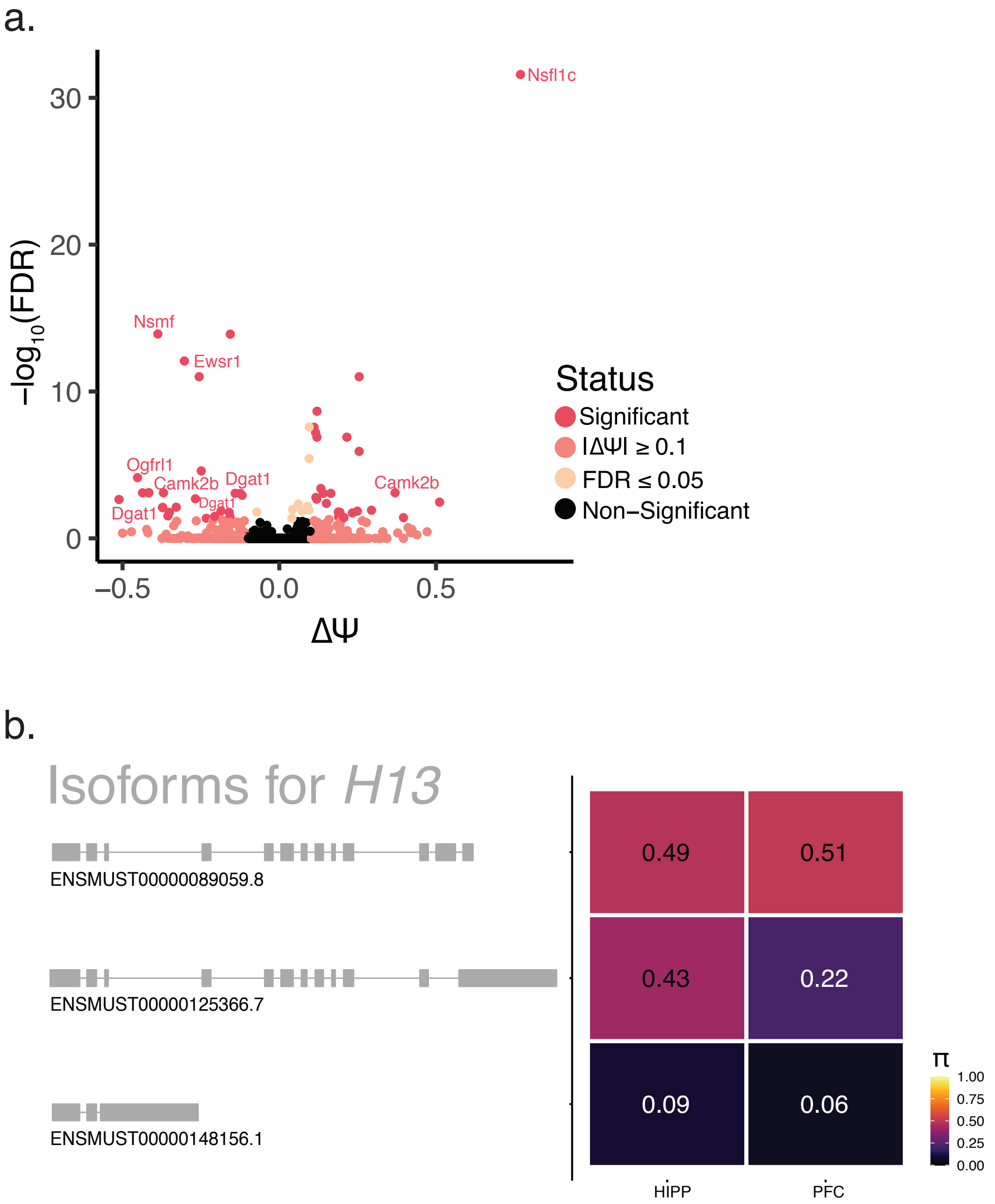


**S5. Gene-wise test for DIE outperforms exon-wise test**

**a.** Volcano plot of bulk HIPP vs. bulk PFC differential exon expression, with the effect size ($\Delta\Psi$) on the X-axis and BY corrected p-value on the Y-axis. Points are colored according to the levels of significance based on FDR and $\Delta\Pi$ value. Genes considered significant (pink) when FDR <= 0.05 and $|\Delta\Psi|$ >= 0.1 **b.** Differential isoform usage missed by exon-based tests because of harsher correction for multiple testing

**Supplemental Figure 6**

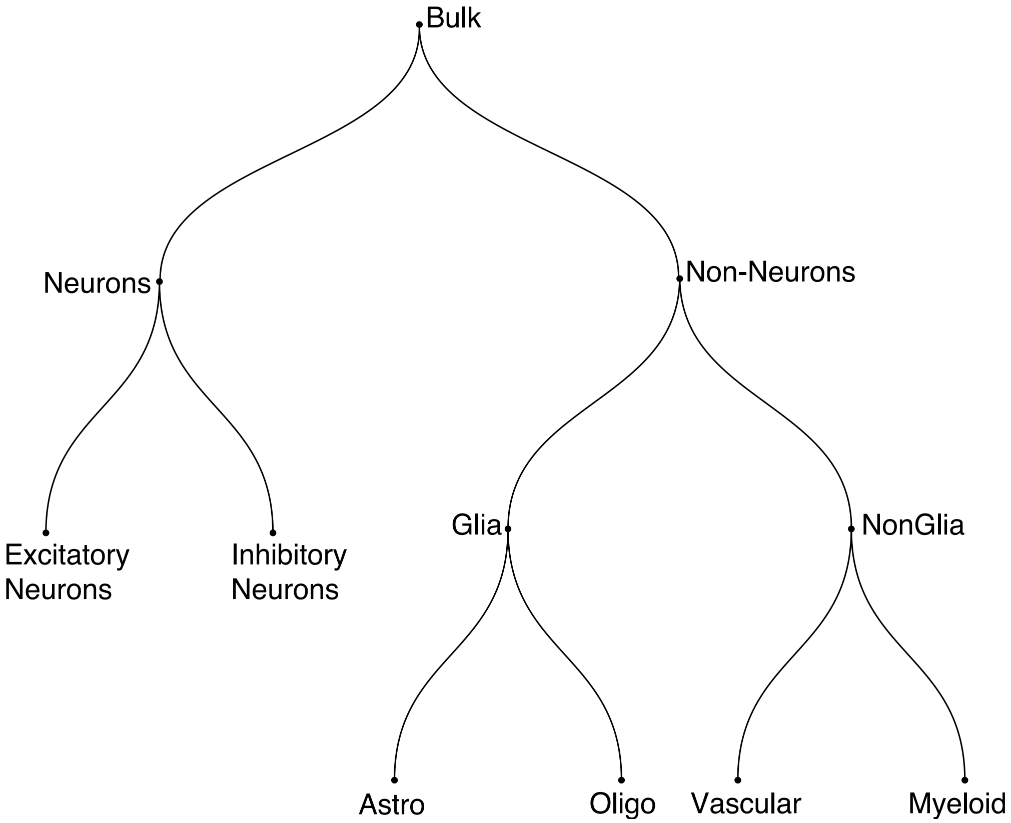


**S6. Cell-type hierarchy based on gene-expression similarities**

**Supplemental Figure 7**

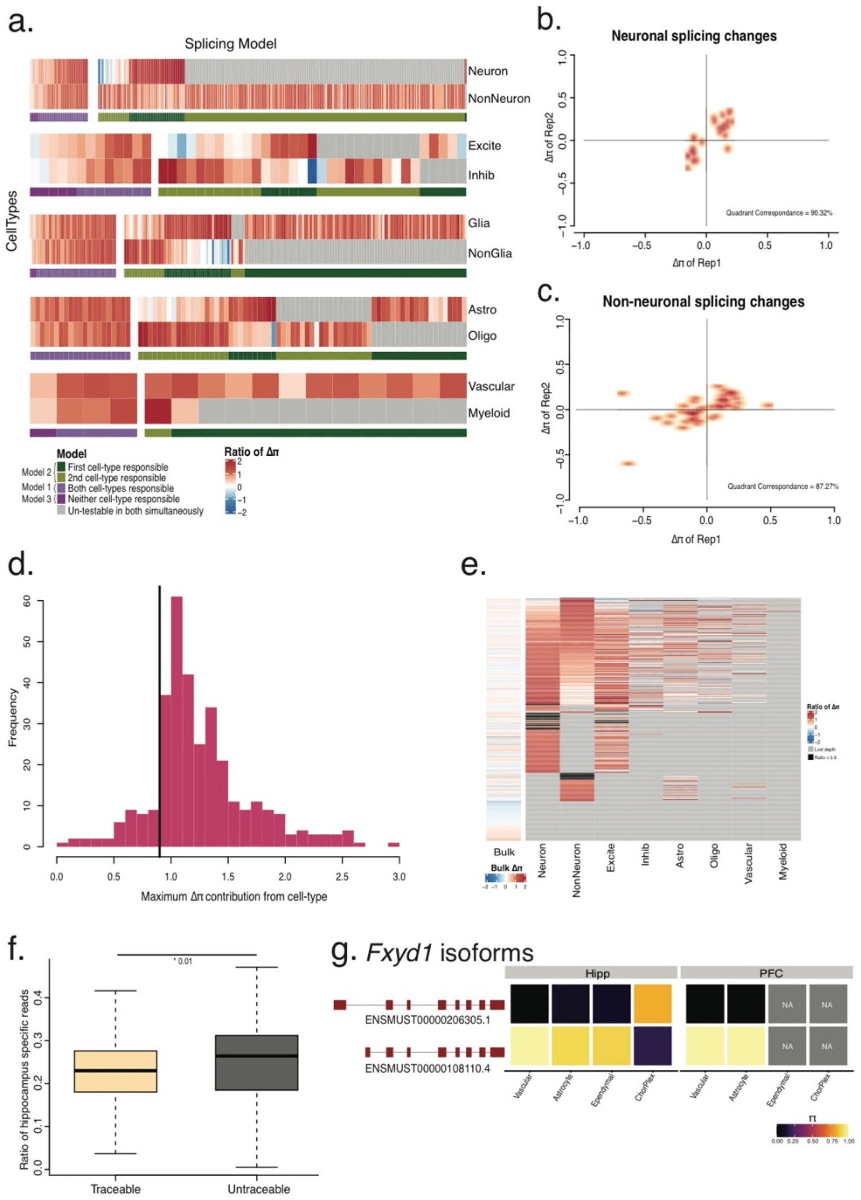


**S7. One cell-type model underlies region-specific splicing differences is replicable**

**a.** Five $gene \times celltype$ heatmaps clustered by the ratio of $\Delta\Pi$ of an individual cell-subtype to a parent cell-type. Each vertical line indicates the ratio of $\Delta\Pi$ for a single gene. Grey lines indicate lack of sufficient depth or lack of expression. Clusters of genes are colored by whether both cell-types show similar relative $\Delta\Pi$ to the parent (purple, Model I, Model III) or whether one cell-type explains most of the splicing changes (Model II) **b.** Scatter plot showing direction of neuronal region-specific splicing changes. X-axis represents $\Delta\Pi$ in neuronal cells of replicate 1. Y-axis represents $\Delta\Pi$ in neuronal cells of replicate 2 **c.** Scatter plot showing direction of non-neuronal region-specific splicing changes. X-axis represents $\Delta\Pi$ in non-neuronal cells of replicate 1. Y-axis represents $\Delta\Pi$ in non-neuronal cells of replicate 2 **d.** Histogram of maximum differential isoform abundance contribution ($\frac{\Delta\pi_{cell-type}}{\Delta\pi_{Bulk}}$) from individual cell groups per gene **e.** Heatmap of $\frac{\Delta\pi_{cell-type}}{\Delta\pi_{Bulk}}$ for 395 region-specific DIE genes for six individual cell-types, and aggregated neurons and non-neurons. Each row is a gene and each column represents the indicated cell-groups. Colors indicate $\frac{\Delta\pi_{cell-type}}{\Delta\pi_{Bulk}}$ except for the bulk column, where colors represent $\Delta\Pi$. Grey indicates untestable in a cell type due to low read depth, while black indicates $\frac{\Delta\pi_{cell-type}}{\Delta\pi_{Bulk}} \leq 0.9$ **f.** Boxplots of ratio of reads originating from hippocampus specific cell-types split by genes where DIE can or cannot be explained by individual cell types **g.** Isoform expression of two major isoforms of *Fxyd1*. Rows indicate isoforms whereas columns indicate cell type split by brain region, colored by $\Pi$ value

**Supplemental Figure 8**

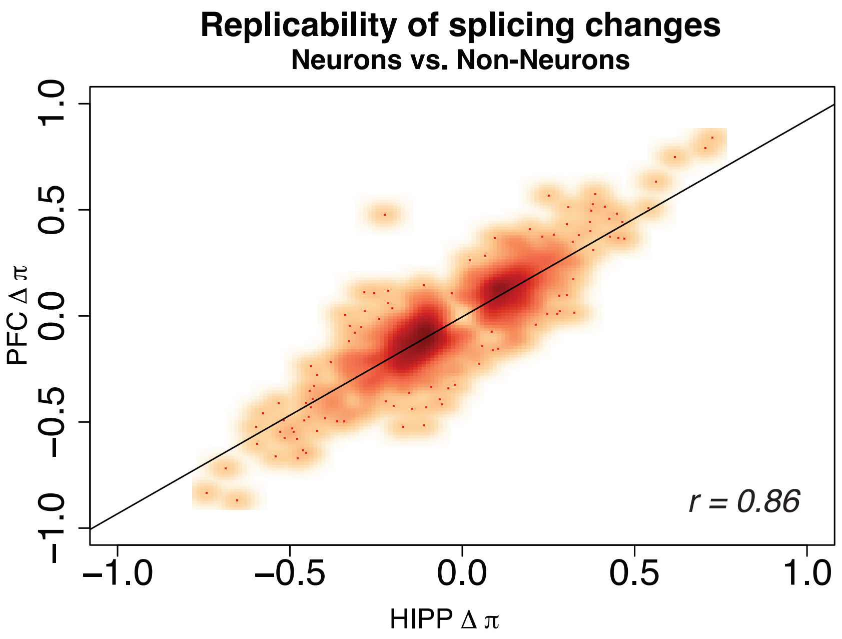


**S8. Regional replicability of cell-type specific splicing changes**

Scatter plot showing the direction of splicing changes between HIPP and PFC for neurons vs. non-neurons. X-axis represents $\Delta\Pi$ of neurons vs non-neurons in hippocampal cells. Y-axis represents $\Delta\Pi$ of neurons vs non-neurons in prefrontal cortex cells

**Supplemental Figure 9**

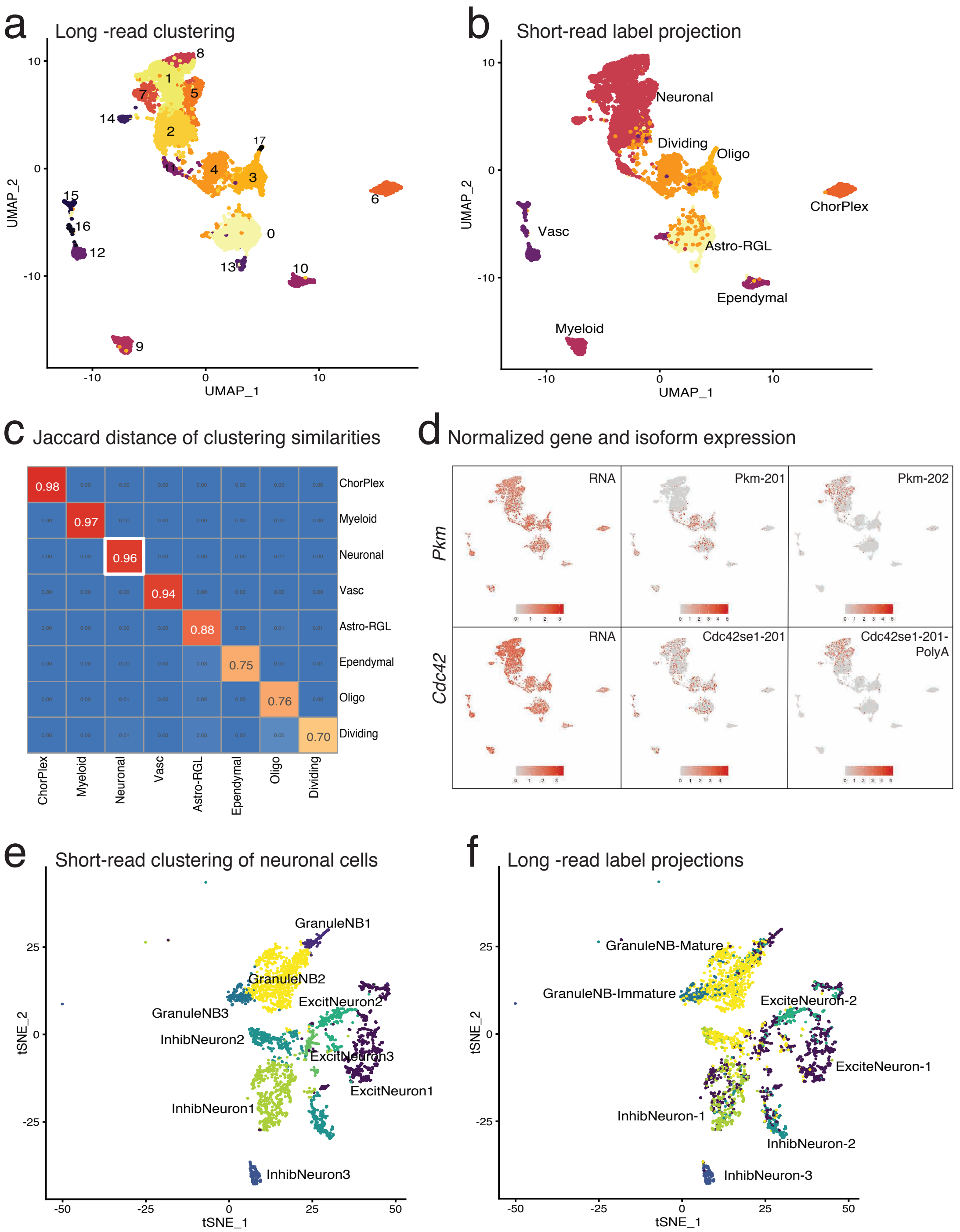


**S9. Clustering on single-cell isoform expression**

**a.** UMAP of clustering based on full-length, hippocampal single-cell transcripts where each point is a cell and color corresponds to various clusters **b.** UMAP from S9A with single-cells colored by cell-types identified by short-read gene expression from Fig1 **c.** Heatmap of jaccard similarity between cell-types identified by clustering on short reads and cell-types identified by clustering on full-length transcripts **d.** Projection of normalized gene and transcript expression for Pkm and Cdc42 on UMAP obtained from clustering on full-length transcripts **e.** Sub-setting neuronal cell-types identified from short-read expression. Clustering based on gene expression similarities between cells, and colored by short-read cell types **f.** Sub-setting neuronal cell-types identified from short-read expression. Clustering based on isoform expression similarities between cells, and colored by cell-types obtained from long-read clustering.

### Supplemental Figure 10

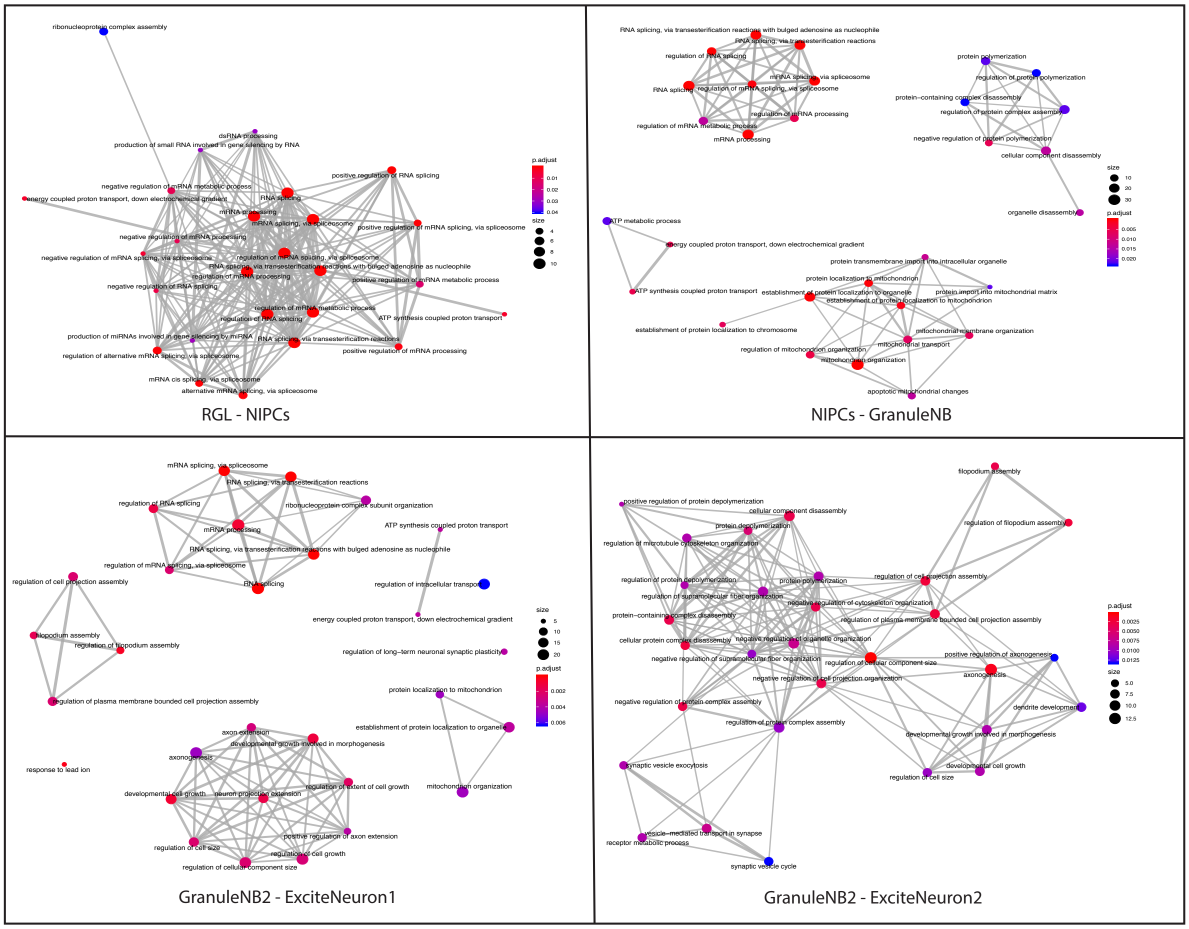


**S10. Network diagrams of DIE genes in the neuronal differentiation trajectory across different comparisons**

Each node represents an enriched gene ontology (GO) term connected by similarity. Color indicates significance level of enrichment, and size indicates the number of significant DIE genes contributing to the GO term.

### Supplemental Figure 11

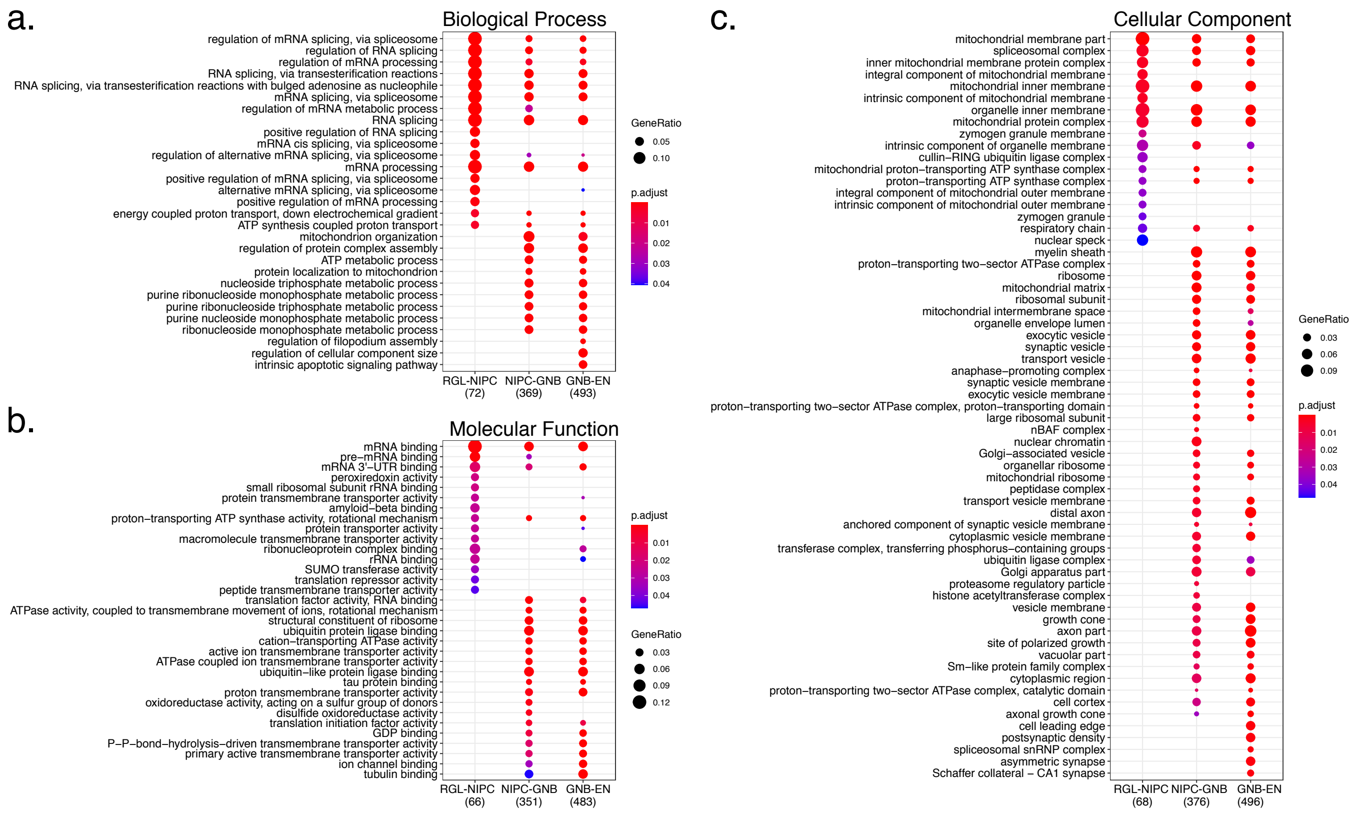


**S11. GO analysis split by category**

**a.** Dotplot of enriched Biological Process GO terms at each step of the neuronal differentiation trajectory in columns. Color indicates significance level of enrichment, and size indicates the number of significant DIE genes contributing to the GO term **b.** Dotplot of enriched Molecular Function GO terms at each step of the neuronal differentiation trajectory in columns. Color indicates significance level of enrichment, and size indicates the number of significant DIE genes contributing to the GO term **c.** Dotplot of enriched Cellular Component GO terms at each step of the neuronal differentiation trajectory in columns. Color indicates significance level of enrichment, and size indicates the number of significant DIE genes contributing to the GO term.

**Supplemental Figure 12**

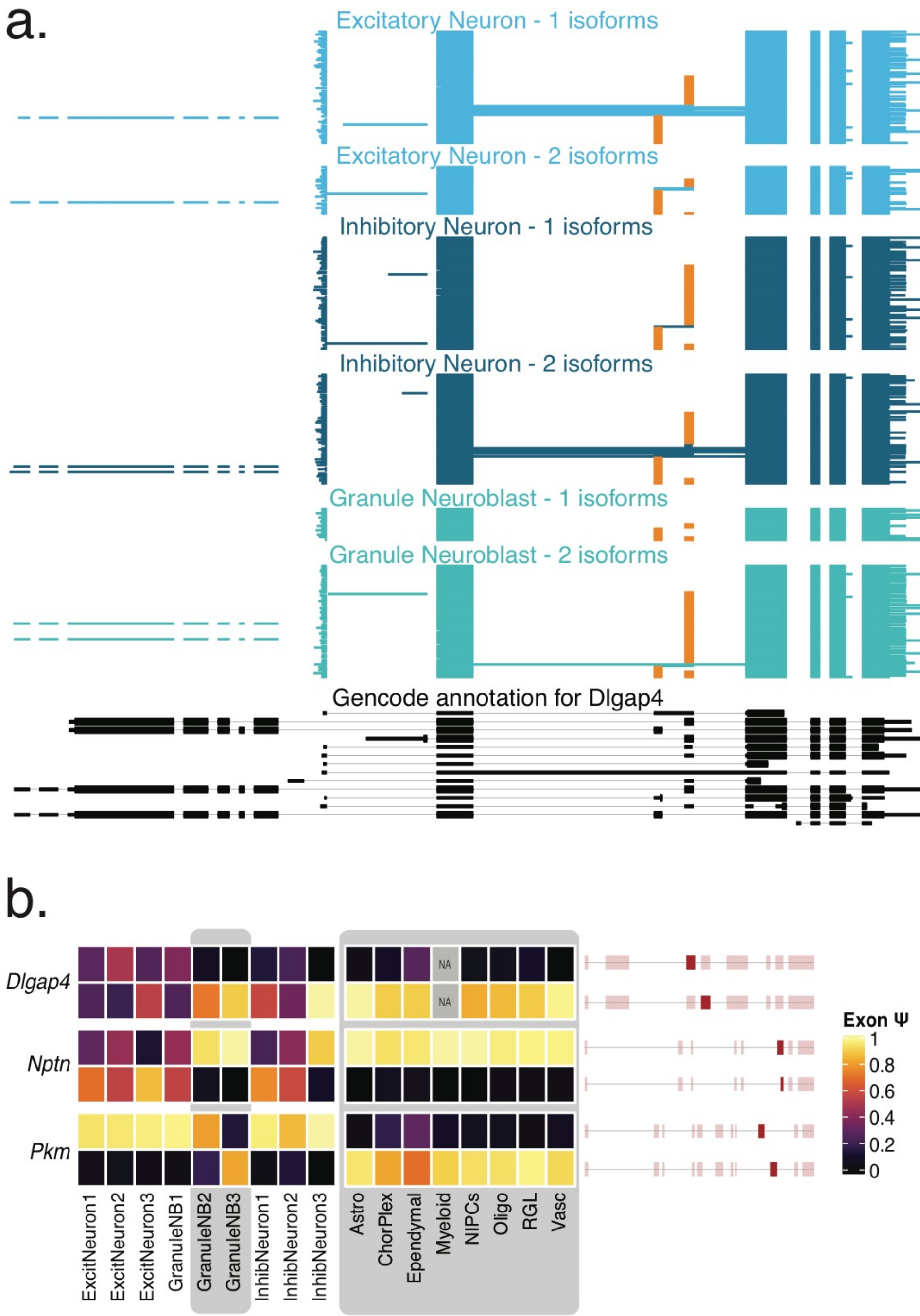


**S12. Cell-type specificity of neurodevelopmentally regulated genes**

**a.** Isoform expression for Dlgap4 gene. Each horizontal line in the plot represents a single transcript colored according to the cell-type it is represented in. Orange exons represent alternative inclusion

**b.** Heatmap of $\Psi$ values for the indicated exons of *Dlgap4, Nptn*, and *Pkm*. Rows indicate exons colored by $\Psi$ values whereas columns indicate the hippocampal cell types they are expressed in. Grey rounded boxes show non-neuronal and immature neuronal cell-types

**Supplemental Figure 13**

a.

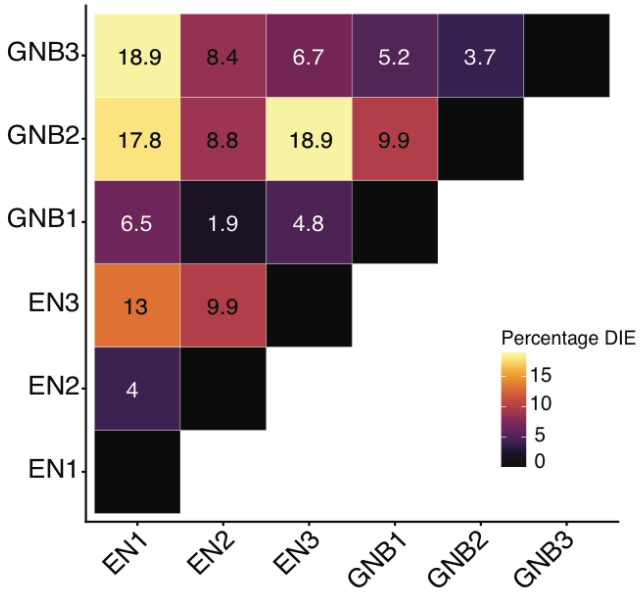


b.

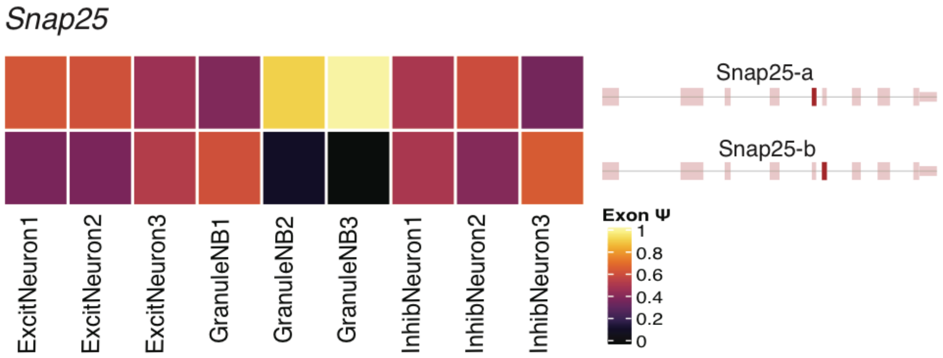


**S13. Cell-type specificity of neurodevelopmentally regulated genes**

**a.** Triangular heatmap showing percentage of DIE genes between all subtypes of GranuleNB (GNB1, GNB2, GNB3) and all subtypes of excitatory neurons (EN1, EN2, EN3)
**b.** Heatmap of $\Psi$ values for the indicated exons of *Snap25*. Rows indicate exons colored by $\Psi$ values whereas columns indicate the neuronal cell types they are expressed in.

**Supplemental Figure 14**

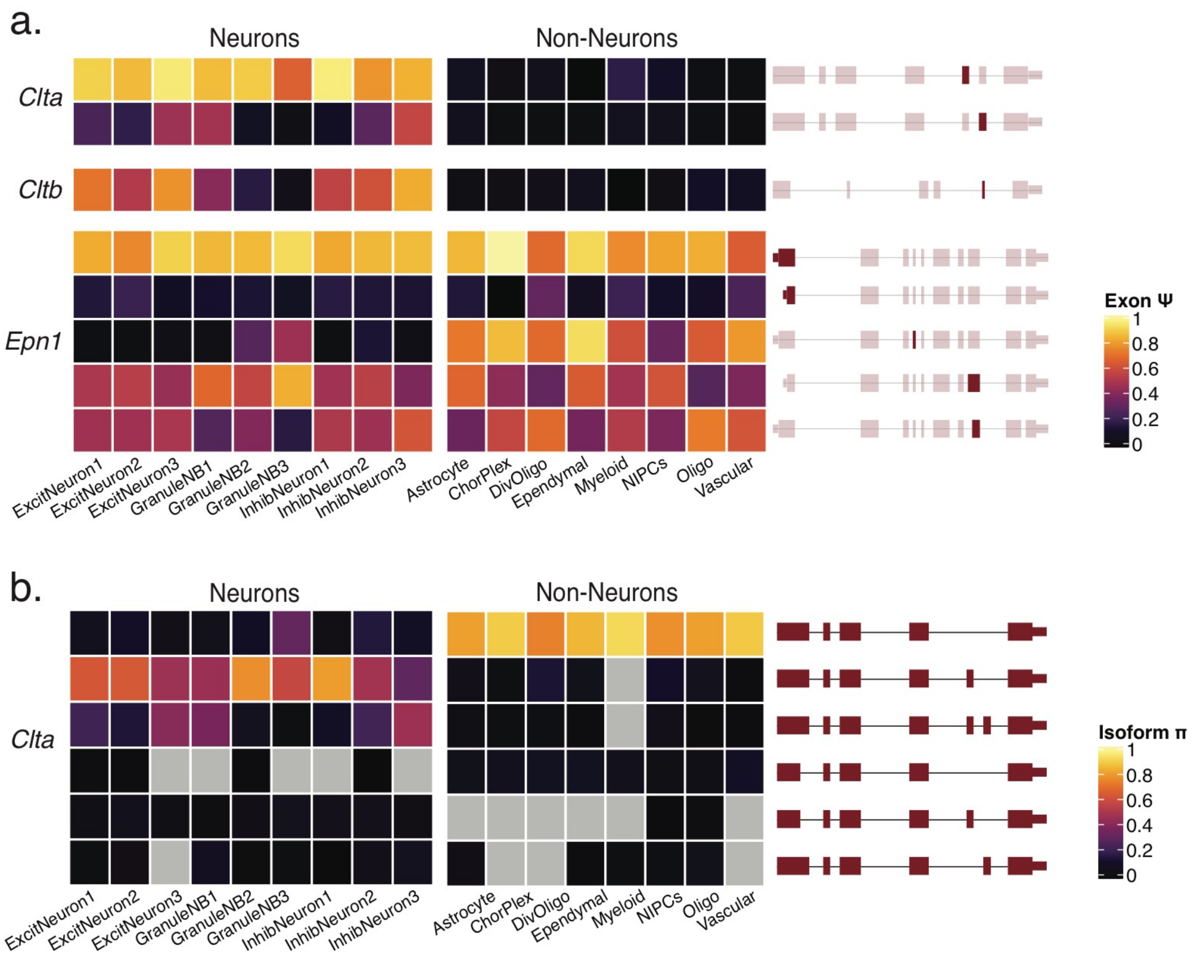


**S14. Cell-type specificity of genes involved in vesicle endocytosis**

**a.** Heatmap of $\Psi$ values for the indicated exons of *Clta, Cltb,* and *Epn*. Rows indicate exons colored by $\Psi$ values whereas columns indicate the hippocampal cell types they are expressed in **b.** Isoform distribution for *Clta*

**Supplemental Figure 15**

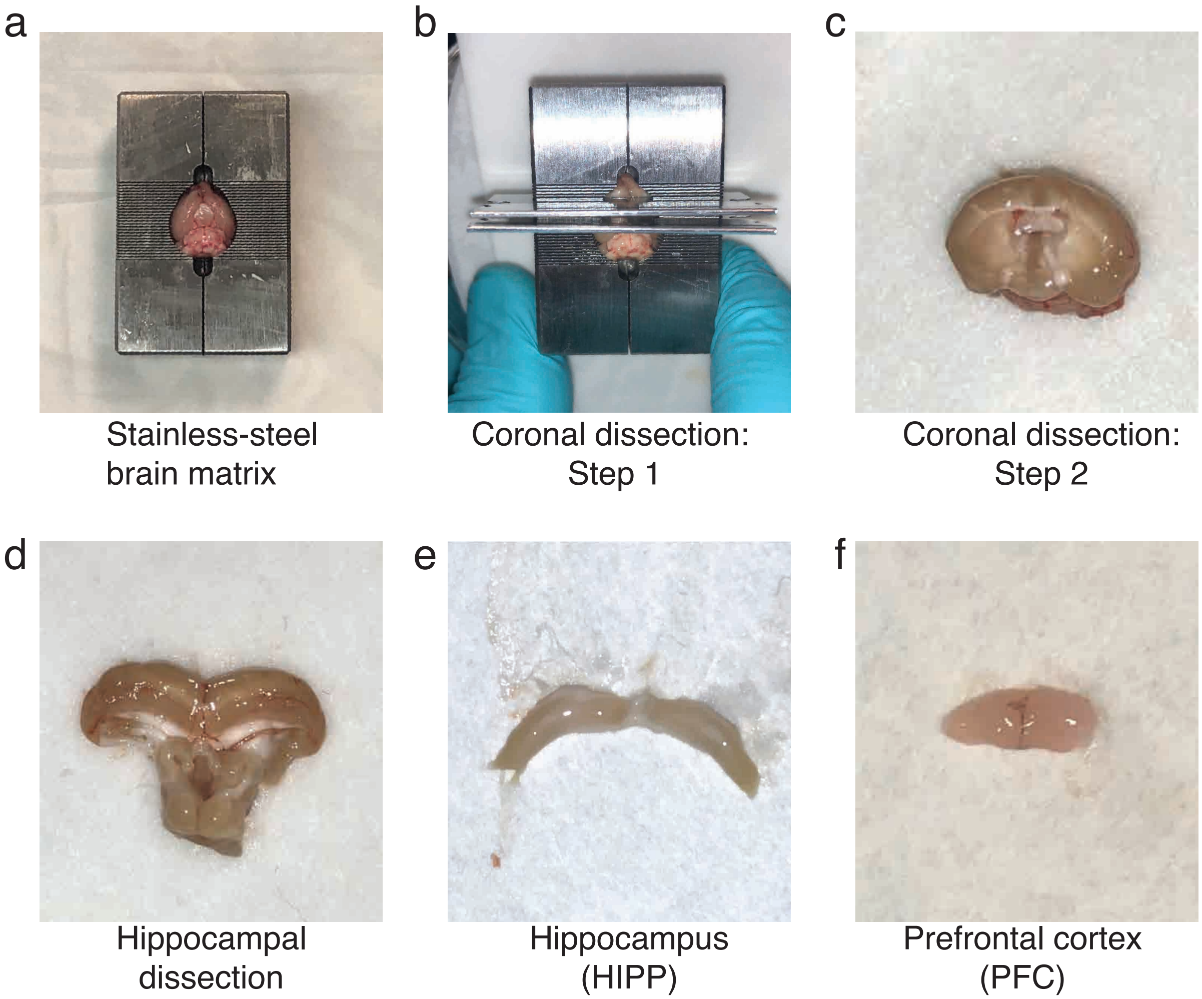


**S15. Steps involved in dissecting the HIPP and PFC**

**a.** Mouse brain placed in stainless steel brain matrix **b.** Step 1 of coronal dissection protocol **c.** Step 2 of coronal dissection protocol **d.** Dissecting out the hippocampal structure **e**. Hippocampus used in experiment **f.** PFC used in experiment

**Supplemental Figure 16**

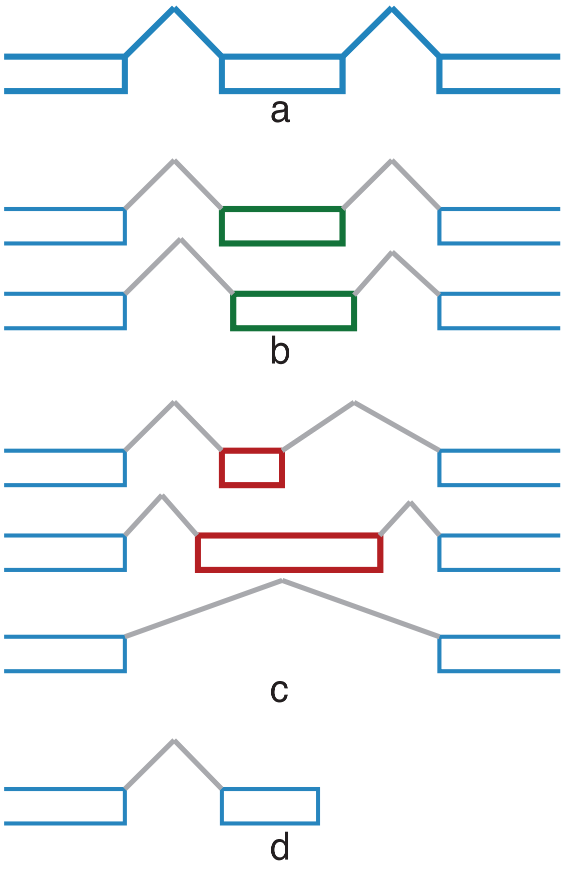


**S16. Schematic showing exon counting**

**a.** Canonical structure of isoform showing annotated internal exon **b.** 3nt difference in splice-site allowed **c.** More than 3nt difference in splice-sites not allowed **d.** Partially covered internal exons not allowed

**Supplemental Table 1**

| Replicate | Region | # reads | Barcoded | Spliced | Barcoded & Spliced | Full-length |
|---|---|---|---|---|---|---|
| Rep1 | HIPP | 38297066 | 15935252 | 14005900 | 6698339 | 4565208 |
| Rep1 | PFC | 10730073 | 3968311 | 4069936 | 1456689 | 1224345 |
| Rep2 | HIPP | 12062107 | 5456753 | 5301119 | 2321631 | 1809583 |
| Rep2 | PFC | 9508271 | 3866334 | 3943532 | 1662952 | 1301187 |
